# Therapeutic and vaccine-induced cross-reactive antibodies with effector function against emerging Omicron variants

**DOI:** 10.1101/2023.01.17.523798

**Authors:** Amin Addetia, Luca Piccoli, James Brett Case, Young-Jun Park, Martina Beltramello, Barbara Guarino, Ha Dang, Dora Pinto, Suzanne M. Scheaffer, Kaitlin Sprouse, Jessica Bassi, Chiara Silacci-Fregni, Francesco Muoio, Marco Dini, Lucia Vincenzetti, Rima Acosta, Daisy Johnson, Sambhavi Subramanian, Christian Saliba, Martina Giurdanella, Gloria Lombardo, Giada Leoni, Katja Culap, Carley McAlister, Anushka Rajesh, Exequiel Dellota, Jiayi Zhou, Nisar Farhat, Dana Bohan, Julia Noack, Florian A. Lempp, Elisabetta Cameroni, Bradley Whitener, Olivier Giannini, Alessandro Ceschi, Paolo Ferrari, Alessandra Franzetti-Pellanda, Maira Biggiogero, Christian Garzoni, Stephanie Zappi, Luca Bernasconi, Min Jeong Kim, Gretja Schnell, Nadine Czudnochowski, Nicholas Franko, Jennifer K. Logue, Courtney Yoshiyama, Cameron Stewart, Helen Chu, Michael A. Schmid, Lisa A. Purcell, Gyorgy Snell, Antonio Lanzavecchia, Michael S. Diamond, Davide Corti, David Veesler

**Affiliations:** Department of Biochemistry, University of Washington, Seattle, WA 98195, USA; Humabs BioMed SA, a subsidiary of Vir Biotechnology, 6500 Bellinzona, Switzerland; Department of Medicine, Washington University School of Medicine, St. Louis, MO, USA; Vir Biotechnology, San Francisco, CA 94158, USA; Faculty of Biomedical Sciences, Università della Svizzera italiana, Lugano, Switzerland; Department of Medicine, Ente Ospedaliero Cantonale, Bellinzona, Switzerland; Clinical Trial Unit, Ente Ospedaliero Cantonale, Lugano, Switzerland; Division of Clinical Pharmacology and Toxicology, Institute of Pharmacological Sciences of Southern Switzerland, Ente Ospedaliero Cantonale, Lugano, Switzerland; Department of Clinical Pharmacology and Toxicology, University Hospital Zurich, Zurich, Switzerland; Division of Nephrology, Ente Ospedaliero Cantonale, Lugano, Switzerland; Clinical School, University of New South Wales, Sydney, New South Wales, Australia; Clinical Research Unit, Clinica Luganese Moncucco, Lugano, Switzerland; Clinic of Internal Medicine and Infectious Diseases, Clinica Luganese Moncucco, Lugano, Switzerland; Division of Nephrology, Cantonal Hospital Aarau, Aarau, Switzerland; Institute of Laboratory Medicine, Cantonal Hospital Aarau, Aarau, Switzerland; Division of Allergy and Infectious Diseases, University of Washington, Seattle, WA 98195, USA; Department of Pathology & Immunology, Washington University School of Medicine, St. Louis, MO, USA; Department of Molecular Microbiology, Washington University School of Medicine, St. Louis, MO, USA; Andrew M. and Jane M. Bursky Center for Human Immunology and Immunotherapy Programs, Washington University School of Medicine, Saint Louis, MO, USA; Center for Vaccines and Immunity to Microbial Pathogens, Washington University School of Medicine, Saint Louis, MO, USA; Howard Hughes Medical Institute, University of Washington, Seattle, WA 98195, USA

## Abstract

Currently circulating SARS-CoV-2 variants acquired convergent mutations at receptor-binding domain (RBD) hot spots^1^. Their impact on viral infection, transmission, and efficacy of vaccines and therapeutics remains poorly understood. Here, we demonstrate that recently emerged BQ.1.1. and XBB.1 variants bind ACE2 with high affinity and promote membrane fusion more efficiently than earlier Omicron variants. Structures of the BQ.1.1 and XBB.1 RBDs bound to human ACE2 and S309 Fab (sotrovimab parent) explain the altered ACE2 recognition and preserved antibody binding through conformational selection. We show that sotrovimab binds avidly to all Omicron variants, promotes Fc-dependent effector functions and protects mice challenged with BQ.1.1, the variant displaying the greatest loss of neutralization. Moreover, in several donors vaccine-elicited plasma antibodies cross-react with and trigger effector functions against Omicron variants despite reduced neutralizing activity. Cross-reactive RBD-directed human memory B cells remained dominant even after two exposures to Omicron spikes, underscoring persistent immune imprinting. Our findings suggest that this previously overlooked class of cross-reactive antibodies, exemplified by S309, may contribute to protection against disease caused by emerging variants through elicitation of effector functions.

---

The emergence of the Omicron variant of concern at the end of 2021 marked a new phase of the COVID-19 pandemic^2^. Omicron lineages accumulated tens of amino acid mutations in their spike (S) glycoprotein that enhanced receptor engagement, altered the preferred internalization route in cells and promoted immune evasion from neutralizing antibodies of unprecedented magnitude^3–15^. As a result, most countries experienced repeated waves of infections in 2021 and 2022, driven by successive Omicron lineages (e.g., BA.1/BA.1.1, BA.2, BA.5), including in individuals who received multiple COVID-19 vaccine doses.

Currently, many different Omicron variant lineages are co-circulating worldwide. Due to convergent evolution, many of these lineages independently acquired identical or similar amino acid mutations at key antigenic sites in the RBD and in the NTD, relative to their presumed BA.2 and BA.5 ancestors^1^. The BA.2.75.2 lineage rose in frequency in multiple countries (e.g., India) and has the following RBD residue mutations relative to BA.2: D339H, R346T, G446S, N460K, F486S and R493Q. CH.1.1 emerged in Asia in November of 2022 and currently accounts for ∼12% of infections in Europe and carries the K444T and L452R RBD residue mutations relative to BA.2.75.2. XBB.1 is an inter-Omicron recombinant lineage, derived from BJ.1 and BA.2.75, and harbors the following RBD residue mutations relative to BA.2: D339H, R346T, L368I, V445P, G446S, N460K, F486S, F490S, R493Q **(Fig. 1a)**. Furthermore, the XBB.1.5 lineage, which contains a proline at position 486 instead of a serine (F486 in Wuhan), is currently rising in the US with ∼85% of cases attributed to the variant as of 25 February 2023. BQ.1 and BQ.1.1 are dominant in several Western countries and account for 56% of all sequenced SARS-CoV-2 genomes in the US (https://covid.cdc.gov/covid-data-tracker/#variant-proportions). BQ1.1 has the following residue mutations relative to BA.5: R346T, K444T and N460K **(Fig. 1a)**. Several of the mutated amino acid positions present in the XBB.1(.5) and BQ.1.1 variant S glycoproteins were previously detected in cryptic lineages through wastewater sequencing^16^ and virtually all of them map to key NTD and RBD antigenic sites.

**Fig. 1.**
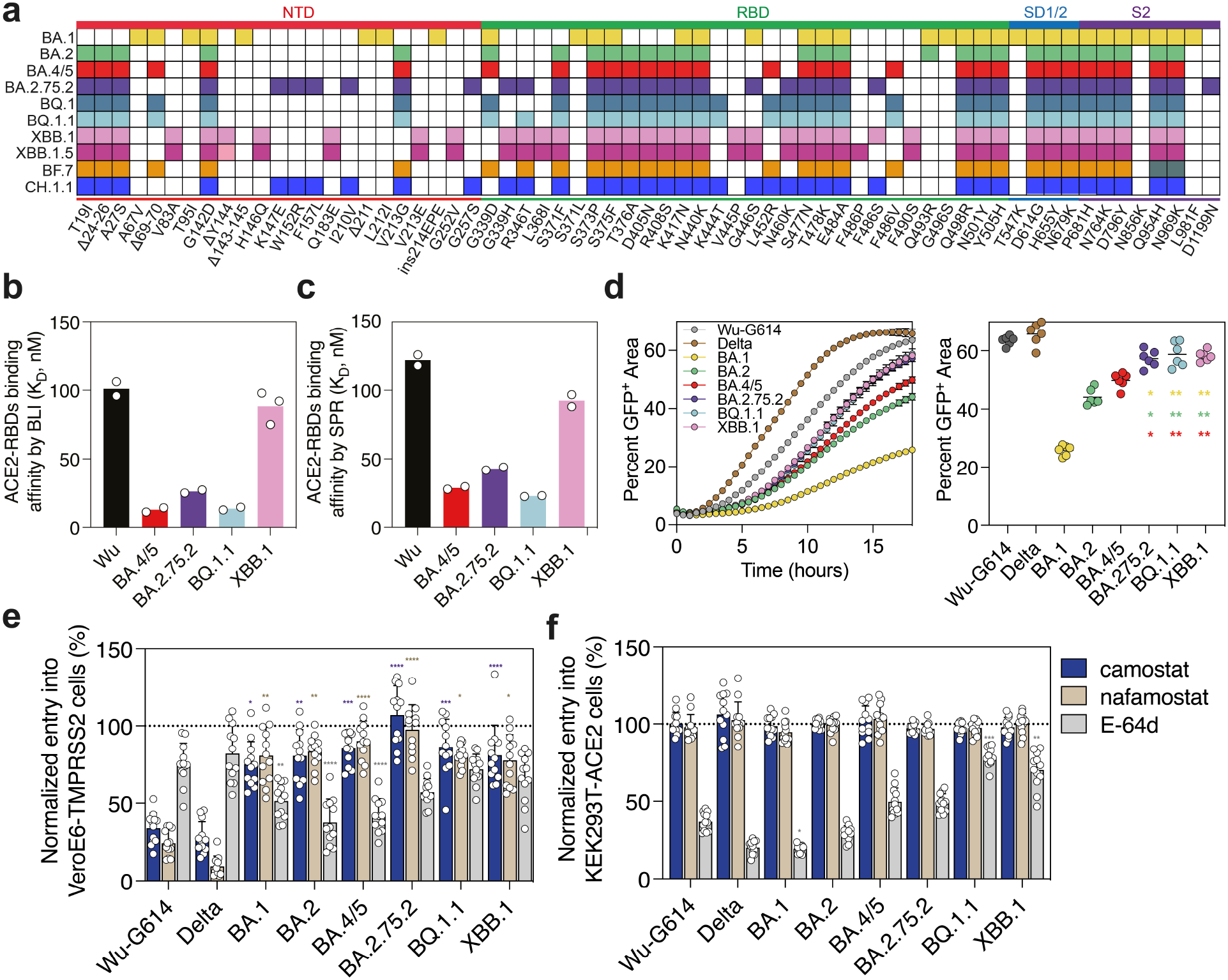
Functional properties of the newly emerged BQ.1.1, XBB.1 and BA.2.75.2 variant S glycoproteins. **a**, Schematic view of S mutations carried by SARS-CoV-2 variants used in this study. **b, c**, Equilibrium dissociation constants (K_D_) measured by biolayer interferometry (b) and surface plasmon resonance (c) for binding of the monomeric human ACE2 ectodomain to the immobilized Wu, BA.4/5, BA.2.75.2, BQ.1.1, or XBB.1 RBDs. **d**, Cell-cell fusion (expressed as the percentage of GFP^+^ area) between cells expressing the Wu-G614, Delta, BA.1, BA.2, BA.4/5, BA.2.75.2, BQ.1.1, or XBB.1 S glycoproteins and VeroE6-TMPRSS2 cells measured over an 18-hour time course experiment using a split GFP system (left panel). The mean magnitude of cell-cell fusion at 18 hours is shown on the right panel. Data are from 6 fields of view from a single experiment and representative of results from two independent biological replicates. Comparisons of fusogenicity mediated by BA.1, BA.2, or BA.4/5 S to BA.2.75.2, BQ.1.1, or and XBB.1 S were completed using the Dunnett’s test with the colors of the asterisks indicating which group the comparison is done with (BA.1: gold; BA.2: green; BA.4/5: red). **P < 0.01. **e, f**, Relative entry of VSV pseudotyped with the Wu-G614, Delta, BA.1, BA.2, BA.4/5, BA.2.75.2, BQ.1.1, or XBB.1 S in VeroE6-TMPRSS2 (e) or HEK293T-ACE2 (f) cells treated with 50 µM of camostat, nafamostat, or E-64d. Normalized entry was calculated based on entry values obtained for VeroE6-TMPRSS2 or HEK293T-ACE2 cells treated with DMSO only for each pseudovirus. Mean normalized entry and standard deviation are presented for each pseudovirus. Twelve technical replicates were performed for each pseudovirus and inhibitor and one experiment representative of two independent biological replicates is shown. Comparison of relative entry values were made between Wu-G614 S VSV pseudovirus and each of the examined SARS-CoV-2 variant S VSV pseudoviruses using the Dunnett’s test. *P < 0.05; **P < 0.01; ***P < 0.001; ****P < 0.0001.

SARS-CoV-2 S recognizes angiotensin-converting enzyme 2 (ACE2) as its main entry receptor, leading to membrane fusion and viral entry^17–20^. RBD-directed antibodies account for most neutralizing activity against vaccine-matched and vaccine-mismatched viruses, whereas the N-terminal domain is mostly targeted by variant-specific neutralizing antibodies^21–34^. Here, we set out to understand how the constellation of mutations in the BQ.1.1, XBB.1(.5) and BA.2.75.2 S variants affect the functional properties of SARS-CoV-2, humoral and memory immunity in humans, and recognition by therapeutic antibodies.

## Functional properties of BQ.1.1, XBB.1 and BA.2.75.2 S

We first determined the binding kinetics and affinity of the monomeric human ACE2 ectodomain to immobilized variant RBDs using biolayer interferometry **(Fig. 1b, Extended Data Fig. 1** and **Supplementary Table 1)**. We measured identical equilibrium dissociation constants for the BQ.1.1 and BA.5 RBDs (K_D_= 12.8 nM and 13.7 nM, respectively), indicating that the additional BQ.1.1 mutations, which map outside the ACE2-binding interface, do not influence receptor engagement **(Fig. 1b, Extended Data Fig. 1** and **Supplementary Table 1)**. The enhanced ACE2 binding affinity of the BA.2.75.2 RBD (K_D_= 26.2 nM), relative to BA.2, results from the R493Q reversion, as G446S has a negligible effect and F486S has a deleterious effect on ACE2 engagement based on mutagenesis and deep-mutational scanning data^35^. We also determined that ACE2 bound to the XBB.1 RBD with an affinity comparable to the Wuhan-Hu-1 (Wu) RBD (K_D_= 88.4 nM and K_D_= 101.1 nM, respectively). As V445P does not change the conformation of the ACE2-bound XBB.1 RBD, relative to BQ.1.1, and none of the three residue substitutions compared to BA.2.75.2 involve side chain-mediated contacts with the host receptor, it is possible that the V445P mutation alters the backbone conformational dynamic of the free XBB.1 RBD and dampens ACE2 binding. We observed a similar ranking of these variant RBDs using surface plasmon resonance (SPR) to determine ACE2 binding affinities **(Fig. 1c, Extended Data Fig. 1** and **Supplementary Table 2)**. Modulation of ACE2 binding resulted from off-rate differences, whereas on-rates remained comparable across all variants tested, in agreement with observations made with previous variants^3,5,35–37^. Collectively these data indicate that BQ.1.1, BA.2.75.2 and BA.5 have comparably high ACE2 binding affinity, suggesting that the BQ.1.1 and BA.2.75.2 viral fitness is not limited by this step of host cell invasion, whereas the lower ACE2-binding affinity of XBB.1 might have limited its spread.

We next compared the kinetics and magnitude of cell-cell fusion promoted by the Wu-G614, Delta, BA.1, BA.2, BA.5, BQ.1.1, BA.2.75.2 and XBB.1 S glycoproteins. This live cell imaging assay uses a split green fluorescent protein (GFP) system with VeroE6 target cells stably expressing TMPRSS2 and GFP β_1-10_ strands and BHK-21 effector cells stably expressing GFP β_11_ strand and transiently transfected with S^3,38^. We observed slower and reduced fusogenicity for the BA.5, BA.2 and BA.1 S glycoproteins compared with Wu-G614 and even more so relative to Delta S^12^ **(Fig. 1d** and **Extended Data Fig. 2)**, in line with previous findings and the lack of syncytia formation observed with authentic viruses^3,7^. BQ.1.1, BA.2.75.2 and XBB.1 S, however, promoted membrane fusion more efficiently than the earlier Omicron variants **(Fig. 1d** and **Extended Data Fig. 2)**, suggesting enhanced fusogenic potential of these recently emerged variants.

We and others previously showed that BA.1, BA.2 and BA.5 had an altered cell entry pathway relative to previous SARS-CoV-2 strains, with Omicron variants entering preferentially through the endosomal entry route (cathepsin-mediated) as opposed to plasma membrane fusion (TMPRSS2-mediated)^6–8,19,39^. To assess the preferred cell entry route of the emerging Omicron variants, we investigated the impact of protease inhibitors on entry of non-replicative vesicular stomatitis virus (VSV) pseudotyped with S glycoproteins into VeroE6-TMPRSS2 cells (enabling both plasma membrane and endosomal entry routes) and HEK293T-ACE2 cells (enabling endosomal entry only). The serine protease (TMPRSS2) inhibitors camostat and nafamostat potently blocked entry of Wu-G614 and Delta S VSV pseudoviruses in VeroE6-TMPRSS2 cells, but had limited effect on any of the Omicron variants, including BQ.1.1, BA.2.75.2 and XBB.1 **(Fig. 1e)**. Reciprocally, the cathepsin B and L inhibitor E64d more significantly inhibited the entry of BA.1, BA.2 and BA.5 S VSV pseudoviruses relative to Wu-G614, whereas no significant difference in entry was measured for Delta, BA.2.75.2, BQ.1.1 or XBB.1 S VSV compared to Wu-G614 in VeroE6-TMPRSS2 cells (**Fig. 1e**). Furthermore, BQ.1.1 and XBB.1 S-mediated entry was also less affected by E-64d than all other variant S evaluated in HEK293T-ACE2 cells **(Fig. 1f** and **Extended Data Fig. 2)**. These findings indicate a possible alteration of protease requirements for BQ.1.1 and XBB.1 S-mediated entry, relative to other Omicron lineages assessed. The inefficient use of TMPRSS2 observed, however, concurs with the identical BQ.1.1, BA.2.75.2 and XBB.1 S_2_ subunit sequences, particularly the presence of the N969K substitution, which was previously identified as the key change accounting for the switch from plasma membrane to endosomal cell entry routes of Omicron variants^6–8^.

## Structural analysis of the impact of BQ.1.1, XBB.1 and BA.2.75.2 RBD mutations on recognition by ACE2 and therapeutic antibodies

To reveal how amino acid substitutions in the BQ.1.1 and XBB.1 RBDs alter receptor recognition and key antigenic sites, we determined cryo-electron microscopy (cryoEM) structures of each RBD bound to the human ACE2 ectodomain and to the Fab fragment of the S309 monoclonal antibody (sotrovimab parent) **(Fig. 2a-b, Extended Data Fig. 3a-e** and **Supplementary Table 3)**. In both structures, the R493Q reversion likely relieves repulsion with ACE2 residue K31 and restores a network of local interactions similar to that made with the Wu RBD^40^ **(Fig. 2c)**. Regarding interactions with therapeutic antibodies, the BQ.1.1 RBD structure shows that the K444T substitution would abrogate salt bridges with the carboxyl side chains of the LY-CoV1404 (bebtelovimab parent) and heavy chain residues D56 and D58 or of the COV2-2130 (cilgavimab parent) heavy chain residue D107 **(Fig 2d, e)**. Moreover, R346T (BQ.1.1/XBB.1) would abrogate a salt bridge with the COV2-2130 heavy chain residue D56; G446S (XBB.1) is expected to reduce COV2-2130 binding sterically and V445P (XBB.1) is expected to reduce binding due to a loss of van der Waals interactions with LY-CoV1404 **(Extended Data Fig. 3f)**. These data explain the decreased LY-CoV1404 binding observed by deep-mutational scanning of yeast-displayed mutant RBDs and the markedly reduced neutralization of LY-CoV1404, COV2-2130 or the COV2-2130/COV2-2196 (Evusheld parent) cocktail against the BQ.1.1 and XBB.1 variants^1,35,41^.

**Fig. 2.**
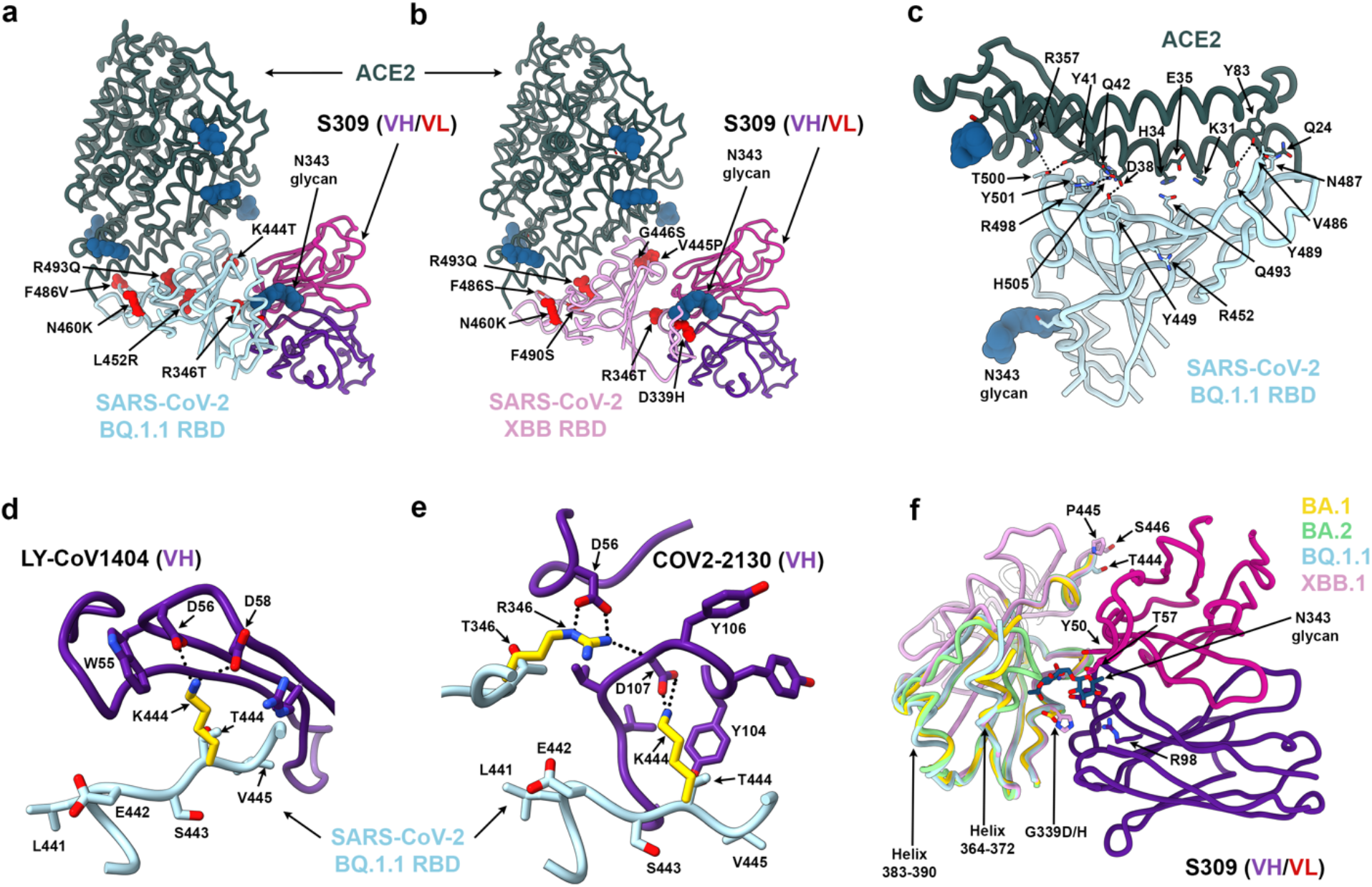
Structural analysis of the BQ.1.1 and XBB.1 RBDs. **a, b**, CryoEM structures of the BQ.1.1 RBD (a, cyan) or the XBB.1 RBD (b, pink) bound to the human ACE2 ectodomain (green) and the S309 Fab fragment (purple/magenta for the heavy/light chains). Amino acid residues mutated relative to Omicron BA.2 are shown as red spheres. **c**, Zoomed-in view of the BQ.1.1 RBD interactions formed with human ACE2 with select amino acid residue side chains shown as sticks. N-linked glycans are shown as dark blue spheres in (a-c). **d, e**, Superimposition of the LY-CoV1404-bound Wu RBD structure (d, purple, PDB 7MMO) or of the COV2-2130-bound Wu RBD structure (e, purple, PDB 7L7E) onto the BQ.1.1 RBD cryoEM structure presented here highlighting the expected disruptions of electrostatic interactions with the monoclonal antibodies resulting from the K444T and the R346T RBD mutations. **f**, RBD-based superimpositions of the S309-bound BA.1 S (gold, PDB 7TLY), apo BA.2 S (green, PDB 7UB0), S309- and ACE2-bound BQ.1.1 (cyan) and XBB.1 (pink) RBD cryoEM structures (this study). S309 is rendered purple and magenta for heavy and light chains, respectively, and the N343 glycan along with select side chains are rendered as sticks. The expected N343 glycan clashes with BA.2 residues N370 and F371 (sticks) are indicated with a red/orange star.

The structures demonstrate that S309 binds to both the BQ.1.1 and XBB.1 RBDs and reveal the molecular basis for accommodation of the H339 residue in the XBB.1 epitope **(Fig. 2f)**. The S309 binding pose is indistinguishable from that observed when bound to the Wu RBD^42^ or to the BA.1 RBD **(Fig. 2f)**. We recently described that the S371F mutation, which is present in BA.2, BA.5, BQ.1.1, XBB.1 and BA.2.75.2, leads to conformational changes of the RBD helix comprising residues 364-372 that are sterically incompatible with the glycan N343 conformation observed in S309-bound spike structures^43^. In the BQ.1.1 structure, helix 364-372 is weakly resolved in the cryoEM map and adopts a conformation similar to that observed in the S309-bound BA.1 structure but distinct from apo BA.2^44^ or apo BA.5 S^45^ structures **(Fig. 2f)**. Residues 368-373 are disordered in the XBB.1 RBD cryoEM map, as is the case for the adjacent residues 380-392 **(Fig. 2f)**. These findings are strongly indicative of conformational frustration of helix 364-372 which is constrained to adopt an energetically disfavored conformation upon S309 binding and could explain the reduced neutralizing activity of this antibody against these variants^1,14,41,46,47^ (**Fig. 3a**).

**Fig. 3.**
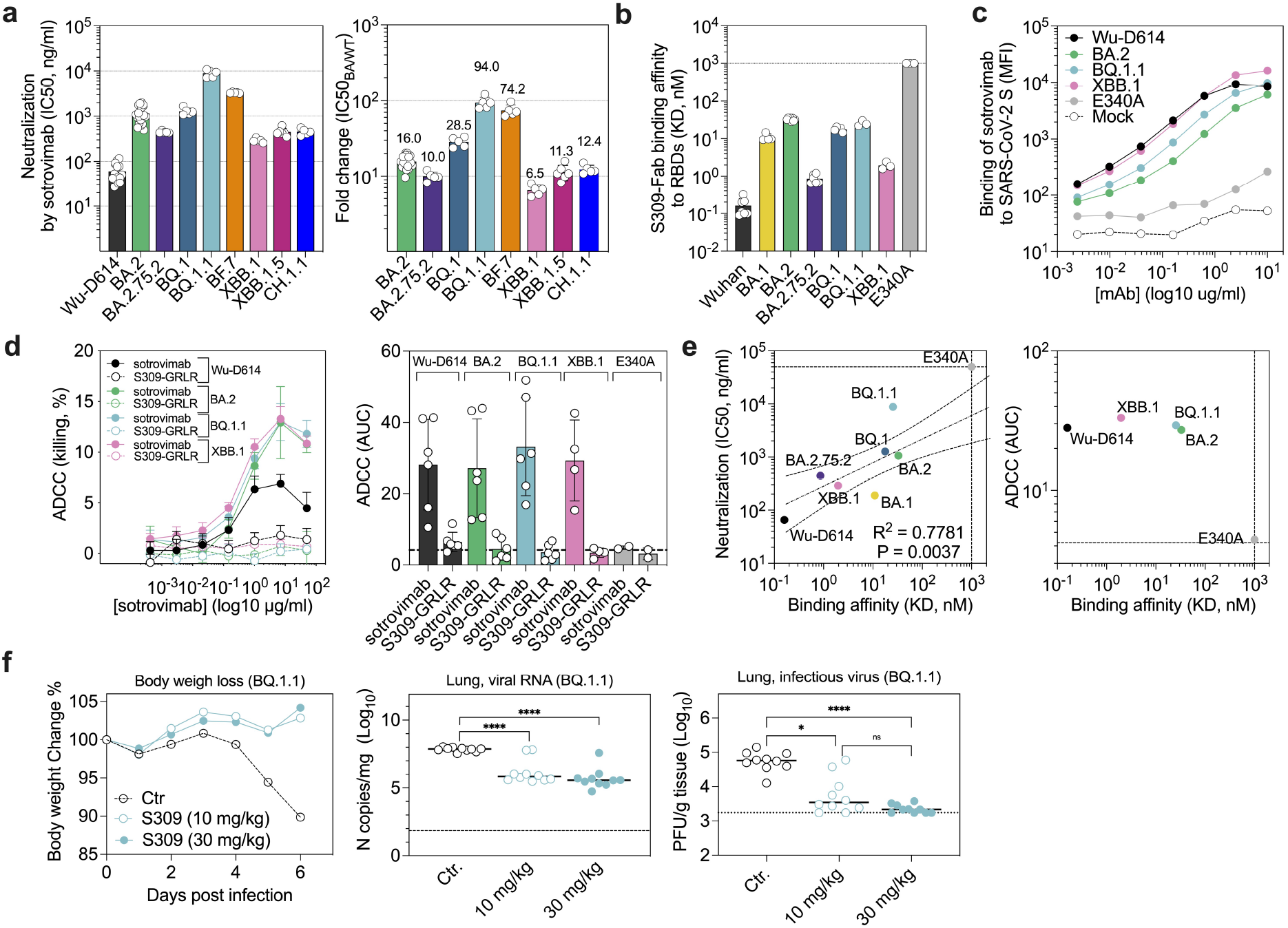
S309 triggers antibody-dependent cell cytotoxicity (ADCC) in vitro and protects mice against BQ.1.1 challenge in vivo. **a**, Sotrovimab-mediated neutralization of Wu-D614, BA.2, BA.2.75.2, BQ.1, BQ.1.1, BF.7, XBB.1, XBB.1.5 and CH.1.1 S VSV pseudoviruses. The sotrovimab neutralizing potency is represented by its IC_50_ (left panel) or fold change relative to neutralization of the Wu-D614 VSV pseudovirus (right panel). Each symbol represents IC_50_ or fold-change values from biological replicates. **b**, Single-cycle kinetics surface plasmon resonance (SPR) analysis of S309 Fab binding to the indicated SARS-CoV-2 RBD variants. Each symbol represents K_D_ values from independent experiments. **c**, Binding of sotrovimab to SARS-CoV-2 S variants transiently expressed at the surface of Expi-CHO cells as determined by flow-cytometry. Similar levels of variant S expression were confirmed by staining with a monoclonal antibody (S2V29) that retains equal and potent neutralizing activity against Wu-D614, BA.2, BQ.1.1, XBB.1 and E340A (**Extended Data Fig. 5a**). **d**, ExpiCHO-S cells transiently transfected with Wu-D614, BA.2, BQ.1.1 or XBB.1 S were incubated with the indicated concentrations of sotrovimab (derivative of S309) or S309-GRLR mAb and mixed with purified NK cells isolated from healthy donors. Cell lysis was determined by a lactate dehydrogenase release assay. Data are presented as mean values ± standard deviations (SD) from one representative donor (left panel). Area under the curve (AUC) analyses from two, four or six NK donors (right panel). **e**, Correlation of sotrovimab Fab binding affinity with neutralizing activity or ADCC. The neutralization IC_50_ values from (a) or the ADCC AUC values from (d) are plotted on the y-axis and the binding affinity to each RBD variant obtained in (b) is plotted on the x-axis. The neutralization data of S309 against BA.1 were adapted from Cameroni et al. Dotted lines indicate the limit of detection. Best-fit lines were calculated using a simple linear regression. Two-tailed Pearson correlation was used to calculate the R^2^ and P values. **f**, Eight-week-old female K18-hACE2 mice received 30 mg/kg of a control isotype-matched monoclonal antibody (anti-West Nile virus hE16), or 10 or 30 mg/kg of S309 (parent of sotrovimab) by intraperitoneal injection one day before intranasal inoculation with 10_4_ FFU of SARS-CoV-2 BQ.1.1. Body weight loss was monitored daily, and tissues were collected at six days after inoculation. Body weight loss (left panel), lung viral RNA determined by RT-qPCR (middle panel), and infectious lung virus titers was measured by plaque assay (right panel) (lines indicate the median; n = 10 mice per group; Kruskal-Wallis ANOVA with Dunn’s post-test between isotype and S309 treatment; ns, not significant; * *P* < 0.05, ****, *P* < 0.0001).

## S309 activates effector functions and protects mice against BQ.1.1 challenge

Based on the cryoEM visualization of S309 binding to the BQ.1.1 and XBB.1 RBDs, we investigated the binding kinetics and affinity of the S309 Fab to the immobilized Wu, BA.1, BA.2 BA.2.75.2, BQ.1, BQ.1.1 and XBB.1 RBDs using SPR **(Fig. 3b)**. The binding affinity (K_D_) of S309 against these variants decreased up to ∼100 fold, primarily as a result of a slower association rate as compared to that against Wu RBD. Little to no dissociation could be observed within the timeframe of our experiment with any RBD evaluated (**Extended Data Fig. 4**). Structural frustration of the RBD helix 364-372, in F371-harboring SARS-CoV-2 Omicron variants, combined with substitutions at position 339 (G339D/H) likely modulate the observed on-rates through S309-mediated backbone conformational and amino acid side chain rotameric selection. We also observed that the sotrovimab IgG (derivative of S309) efficiently cross-reacted with full-length, cell-surface expressed BQ.1.1 and XBB.1 S trimers, to levels comparable or greater than those observed for BA.2 S, but not with the E340A S (negative control^48^) escape mutant **(Fig. 3c** and **Extended Data Fig. 5a)**. These data show that sotrovimab IgG binds avidly to all currently dominant Omicron SARS-CoV-2 variants. However, the neutralizing potency of sotrovimab varied widely against these variants, ranging from a 6.5-fold loss against XBB.1 up to a 94-fold loss against BQ.1.1 relative to Wu **(Fig. 3a** and **Extended Data Fig. 6a-b)**, in line with recent reports^1,14,41,46,47^.

Given the reduced neutralization potency of sotrovimab against these variants despite retaining binding avidity, we evaluated the ability of this therapeutic antibody to activate antibody-dependent cell cytotoxicity (ADCC) using primary natural killer (NK) effector cells and CHO target cells expressing SARS-CoV-2 S of the different Omicron variants at their surface. Sotrovimab efficiently promoted ADCC of cells expressing Wu-D614, BA.2, BQ.1.1 or XBB.1 S in a concentration- and Fc-dependent manner **(Fig. 3d** and **Extended Data Fig. 5b)**. Although we observed a linear relationship between the S309 Fab binding affinity and in vitro neutralization potency, this correlation was not apparent with respect to the magnitude of ADCC, which was equivalent against all variants in spite of up to 100-fold differences in binding affinity **(Fig. 3e)**.

To better understand the physiological relevance of these findings, we administered S309 to mice which were subsequently challenged with BQ.1.1, the variant associated with the greatest loss of in vitro neutralizing activity (i.e., 94-fold loss relative to Wu-D614 S pseudovirus). S309 was given as prophylaxis at 10 or 30 mg/kg doses to K18/hACE2 mice one day prior to challenge with SARS-CoV-2 BQ.1.1. Notably, S309 completely protected K18-hACE2 transgenic mice from weight loss at both doses, whereas animals receiving a control antibody lost greater than 10% of body weight by day 6 **(Fig. 3f)**. Furthermore, S309 administration reduced both viral RNA and infectious virus titers in the lung in a statistically significant manner at both 10 and 30 mg/kg doses relative to control mice **(Fig. 3f)**. These data indicate that, despite its marked reduction of in vitro neutralizing activity, S309 can protect mice from BQ.1.1 challenge (consistent with a recent pre-print on BQ.1.1-challenged hamsters^49^).

## Bivalent mRNA vaccines enhance neutralizing antibody responses against emerging SARS-CoV-2 variants

To assess the impact of the BQ1.1, XBB.1 and BA.2.75.2 S mutations on vaccine-elicited antibody responses, we quantified plasma neutralizing activity using VSV pseudotyped with Wu-G614, BA.1, BA.5, BF.7, BQ.1.1, XBB.1 or BA.2.75.2 S. We compared plasma from eight cohorts of individuals obtained 15-30 days post vaccination or PCR-confirmed breakthrough (BT) infection corresponding to subjects: (i) mRNA vaccinated four times with the prototype vaccine encoding Wuhan-1 spike and without known infection (“Wu_4_ vaccinated”); (ii) mRNA vaccinated four times without known infection, the last dose being the Wu/BA.5 bivalent booster (“Wu/BA.5 bivalent vaccinated”); (iii) previously infected in 2020 (with a WA1/2020–like SARS-CoV-2 strain) and then mRNA vaccinated four to five times, the last dose being Wu/BA.5 mRNA bivalent booster (“pre-Omicron infected-Wu/BA.5 bivalent vaccinated”); (iv) mRNA vaccinated before experiencing an Omicron BA.1, BA.2, BA.2.12.1 or BA.5 BT infection and vaccinated again with the Wu/BA.5 mRNA bivalent booster (“Omicron BT-Wu/BA.5 bivalent vaccinated”). As an alternative to the Wu/BA.5 bivalent mRNA booster, Switzerland and a few other countries, offer a Wu/BA.1 bivalent mRNA booster. We therefore analyzed neutralization in additional cohorts: (v) mRNA vaccinated three times with the Wu monovalent vaccine without known infection (“Wu_3_ vaccinated”); (vi) mRNA vaccinated three times with the Wu monovalent vaccine after pre-Omicron BT infection (“pre-Omicron infected-Wu vaccinated”); (vii) mRNA vaccinated four times without known infection, the last dose being the Wu/BA.1 bivalent booster (“Wu/BA.1 bivalent vaccinated”); (viii) mRNA vaccinated four times with a BA.1 or a BA.2 BT infection, the last dose being the Wu/BA.1 bivalent booster (“Omicron BT-Wu/BA.1 bivalent vaccinated”). In addition, we analyzed plasma samples obtained from kidney transplant recipients (KTR) vaccinated with four Wu monovalent doses with or without pre-Omicron infection.

Wu/BA.5 or Wu/BA.1 bivalent mRNA vaccination elicited comparable neutralizing antibody titers against Wu-G614 S VSV with that observed in matched Wu vaccinated cohorts but higher neutralization of BA.1 S and BA.5 S VSV pseudoviruses **(Fig. 4 a, b)**. Cohorts that received the Wu/BA.5 or the Wu/BA.1 bivalent mRNA vaccines had detectable neutralizing activity against the vaccine-mismatched XBB.1, BA.2.75.2 and BQ.1.1 S VSV pseudoviruses, irrespective of prior infection status, whereas little to no neutralization of these variants was detected for the Wu-only vaccinated subjects **(Fig. 4 a, b)**. We did not detect plasma neutralizing activity against circulating Omicron variants in subjects that were vaccinated four times with the Wu monovalent booster but were immunosuppressed following kidney transplantation, underscoring the difficulties associated with protecting these at-risk populations (**Extended Data Fig. 7**). Overall, these data suggest that bivalent (Wu/BA.1 or Wu/BA.5) mRNA vaccination elicits more potent and broader antibody responses against vaccine-matched and mismatched Omicron variants than Wu mRNA vaccination in healthy subjects.

**Fig. 4.**
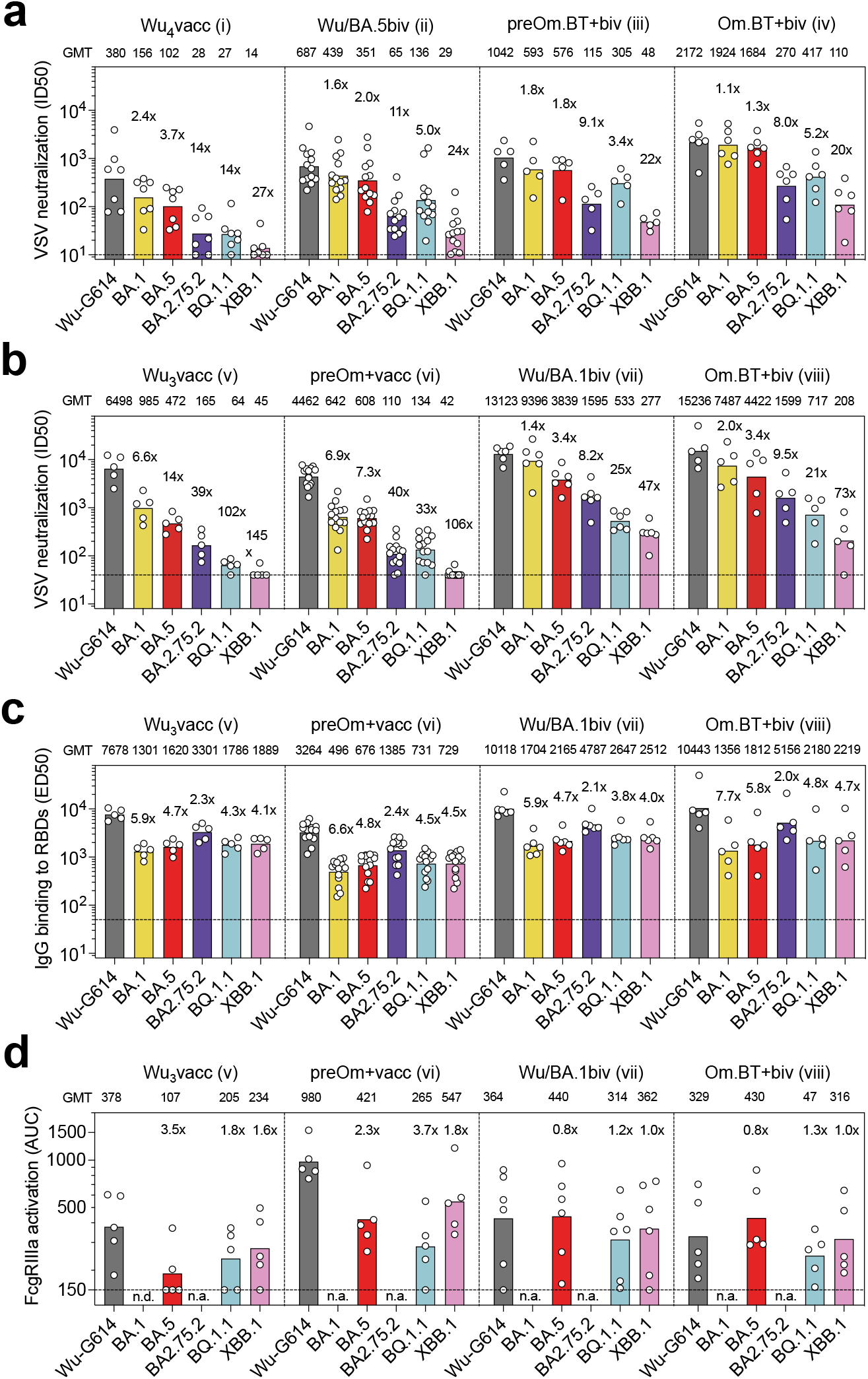
Neutralization, binding and Fc-dependent effector functions of vaccine-elicited plasma antibodies against the BA.2.75.2, BQ.1.1 and XBB.1 variants. **a, b**, Neutralization of SARS-CoV-2 pseudotyped VSV carrying Wu-G614, Omicron BA.1, BA.5, BA.2.75.2, BQ.1.1 and XBB.1 S by plasma samples of cohorts i-iv (a) and v-viii (b). Plasma neutralizing titers expressed as ID_50_s from n = 2 biological (a) and technical (b) replicates are shown. Bars and values on top represent geometric mean ID_50_ titers (GMT). Fold-loss of neutralization against Omicron variants as compared to Wu-G614 is shown above each corresponding bar. Horizontal dashed lines indicate the limit of detection in these assays (ID_50_ = 10 in panel a and 40 in panel b). **c**, Binding of plasma IgG antibodies to SARS-CoV-2 RBDs from Wu, Omicron BA.1, BA.5, BA.2.75.2, BQ.1.1 and XBB.1 variants as measured by ELISA. Shown are ED_50_ values from n = 2 technical replicates of samples from cohorts v-viii. Bars represent geometric mean ED_50_ binding titers (GMT). Fold-loss of binding titers to Omicron variants as compared to Wu is shown above each corresponding bar. Horizontal dashed line indicates the limit of detection (ED_50_ = 50). **d**, Activation of Fc*γ*RIIIa (V158 allele) measured using Jurkat reporter cells and Wu-G614, BA.5, BQ.1.1 and XBB.1 SARS-CoV-2 S glycoprotein-expressing ExpiCHO as target cells. Shown are AUC values from one experiment with plasma samples from cohorts v-viii (n=5 donors for cohort v, n=5 for cohort vi, n=6 for cohort vii and n=5 for cohort viii). Bars and values on top represent geometric mean AUC titers (GMT). Fold-change of activation with Omicron variants as compared to Wu-G614 is shown above each corresponding bar. The horizontal dashed line indicates the limit of detection (AUC = 150). n.a., not assayed. Demographics of cohorts are summarized in **Supplementary Table 5**.

## Plasma antibodies maintain binding and Fc-mediated effector function against emerging Omicron variants

Next, we investigated binding and Fc-mediated effector function of plasma antibodies from cohorts v to viii. The progressive reduction of neutralizing antibody titers against currently dominant Omicron variants overtime was not paralleled by a similar decrease in IgG binding titers to matched RBDs (**Fig. 4c**), as binding titers remained comparable across all Omicron variants. This finding is consistent with data on pre-Omicron variants^50^ and with the notion that SARS-CoV-2 is evolving primarily to escape from neutralizing antibodies that exert the greatest selective pressure.

Fc-mediated effector function of plasma antibodies was evaluated by measuring activation of the Fc*γ*RIIIa (V158 allele) as a surrogate of ADCC using Jurkat reporter cells and Wu-G614, BA.5, BQ.1.1 and XBB.1 SARS-CoV-2 S-expressing ExpiCHO as target cells (**Fig. 4d** and **Extended Data Fig. 8**). Each cohort exhibited high individual variability in the ability of plasma to trigger activation of Fc*γ*RIIIa, with some donors lacking measurable activity despite the presence of high titers of RBD-binding antibodies (**Fig. 4c**). Samples that scored positive for Fc*γ*RIIIa triggering against Wu retained activity against BA.5, BQ.1.1 and XBB.1 S variants. These results suggest that antibodies capable of triggering Fc effector functions vary widely amongst individuals, but, when present, these antibodies are broadly reactive against Omicron variants.

## Cross-reactive RBD-specific memory B cells dominate bivalent vaccine-elicited responses

We next compared memory B cell (MBC) populations in the Wu_4_ vaccinated, Wu/BA.5 bivalent vaccinated (including one individual [31H] who received the Janssen COVID-19 vaccine as their first vaccine series), pre-Omicron infected-Wu/BA.5 bivalent vaccinated, and Omicron BT-Wu/BA.5 bivalent vaccinated subjects (cohorts i-iv) by enumerating the frequency of Wu RBD-specific, Omicron RBD-specific (pool of BA.1/BA.2/BA.5), and cross-reactive MBCs by flow cytometry. Individuals who were exposed only to Wu S through vaccination (Wu_4_ vaccinated) had the highest frequency of Wu RBD-specific MBCs (mean frequency: 25.8%) and the lowest frequency of cross-reactive MBCs (mean frequency: 71.1%) of the four cohorts examined **(Fig. 5a, b** and **Extended Data Fig. 9)**. Individuals who were exposed only once to Omicron S through vaccination had few Omicron RBD-specific MBCs (mean frequency: 5.1%), regardless of whether they had experienced a pre-Omicron SARS-CoV-2 infection (Wu/BA.5 bivalent and pre-Omicron bivalent). Most RBD-specific MBCs in these cohorts cross-reacted with the Wu RBD and the Omicron RBD pool (BA.1/BA.2/BA.5), with uninfected individuals having similar frequencies of cross-reactive MBCs (mean: 77.4%) when compared to individuals who had experienced a pre-Omicron infection (mean frequency: 77.1%) **(Fig. 5a, b** and **Extended Data Fig. 9)**. These data are consistent with previous analyses of MBC populations in Wu vaccinated individuals who had experienced an Omicron BT infection and suggest that immunological imprinting limits the development of new Omicron-specific MBCs, although there is efficient recall of cross-reactive MBCs after a single exposure to Omicron S^1,43,51^. Although Omicron BT-Wu/BA.5 bivalent vaccinated subjects had two exposures to an Omicron S (one through infection and one through vaccination), they still had relatively few Omicron-specific MBCs (mean: 5.3%) similar to those individuals who received only the bivalent booster. The Wu/Omicron (BA.1/BA.2/BA.5) RBD pool cross-reactive MBCs were further enriched (mean frequency: 81.5%) for this cohort compared to the Wu/BA.5 bivalent vaccinated cohort **(Fig. 5a, b** and **Extended Data Fig. 9)**.

**Figure. 5.**
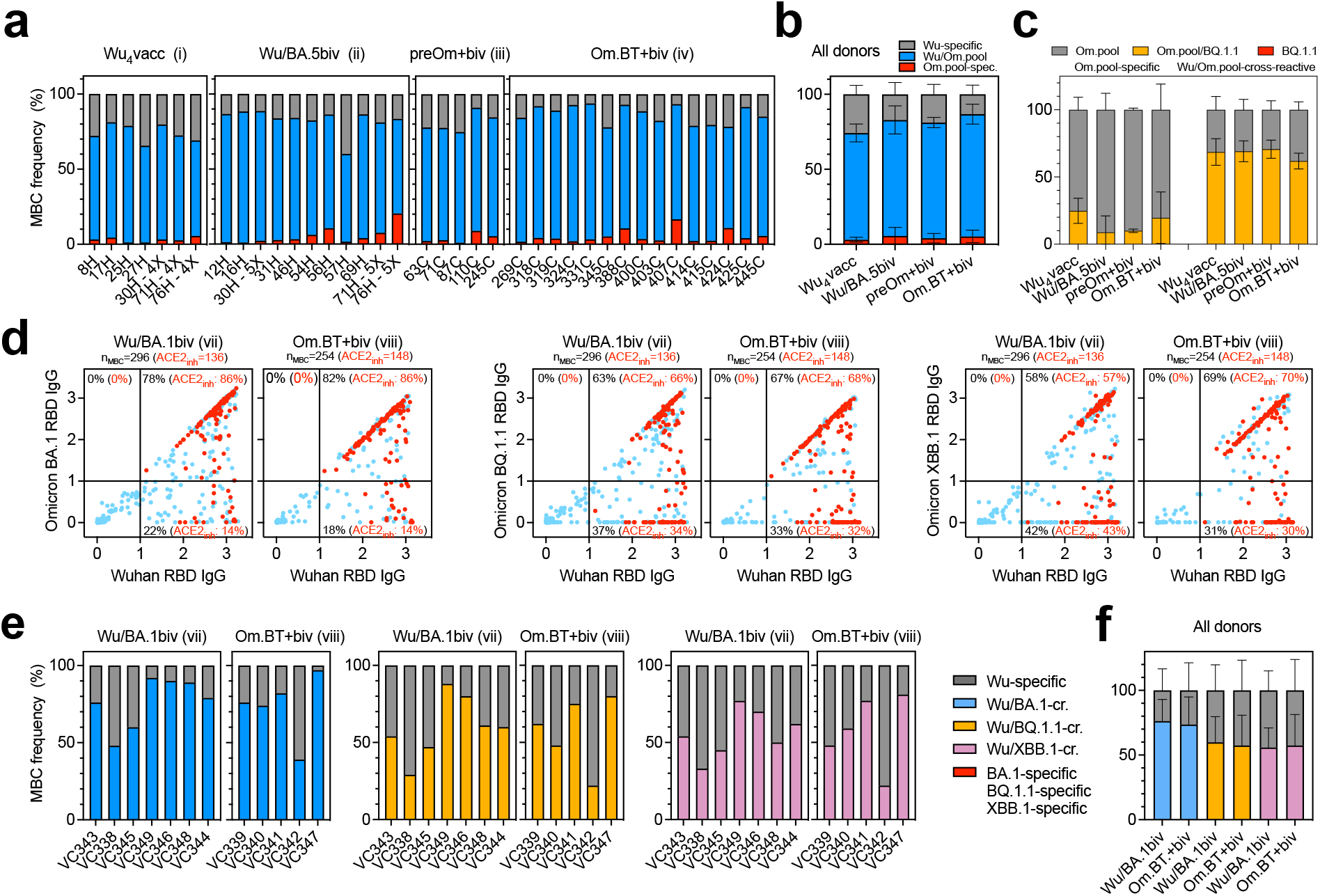
Cross-reactivity of vaccine-elicited SARS-CoV-2 RBD-specific MBCs. **a, b**, Frequency of Wu RBD-specific (grey), Omicron (BA.1/BA.2/BA.5) RBD pool-specific (red) and Wu/Omicron RBD pool-cross-reactive (blue) MBCs from donors of cohorts i-iv, as measured by flow cytometry. Individual frequencies and mean ± SD are shown in panels a and b, respectively. **c**, Analysis of cross-reactivity with the BQ.1.1 RBD of Omicron RBD pool-specific (red bars of panel b) and Wu/Omicron RBD-cross-reactive (blue bars of panel b) MBCs. Om.pool, MBCs recognizing the Omicron (BA.1/BA.2/BA.5) RBD pool. **d**, Cumulative cross-reactivity of IgGs secreted from in vitro stimulated MBCs between the Wu RBD and the Omicron BA.1, BQ.1.1 or XBB.1 RBDs as measured by ELISA. Data represent average OD values with blank subtracted from n = 2 replicates of MBC cultures analyzed from donors of cohorts vii and viii. RBD-specific IgGs showing inhibition of binding to ACE2 are depicted in red. Number of total and ACE2-inhibiting (ACE2_inh_) RBD-directed IgG positive cultures are indicated on top of each graph. Percentages of Wu-specific, Omicron-specific and Wu/Omicron-cross-reactive IgG positive cultures are indicated within each quadrant. **e, f**, Individual frequencies and mean frequencies ± SD of Wu RBD-specific (grey), Omicron-specific (red) and Wu/Omicron RBD cross-reactive (blue for BA.1, yellow for BQ.1.1 and purple for XBB.1) IgG positive cultures from donors of cohorts vii and viii are shown in panels **e** and **f**, respectively.

To assess the ability of MBCs to recognize the currently circulating Omicron variants, we examined whether MBCs recognizing the Omicron RBD pool (including those that are cross-reactive with the Wu RBD and those that are not) could bind the BQ.1.1 RBD (**Fig. 5c** and **Extended Data Fig. 10a, b**). Most Wu/Omicron RBD pool cross-reactive memory B cells recognized the BQ.1.1 RBD (mean frequency: 66.3%), whereas only a small fraction of MBCs binding to the Omicron RBD pool and not the Wu RBD also recognized the BQ.1.1 RBD (mean frequency: 16.9%), regardless of infection and vaccination status. These Omicron-specific MBCs were likely elicited de novo upon Omicron S exposure (through infection and/or vaccination) and their breadth towards currently dominant Omicron variants may improve over time through affinity maturation, similar to the maturation of the antibody response observed after infection with the SARS-CoV-2 Wu or Washington-1 strain or vaccination^30,52–59^.

We next determined if the BA.5 versus BA.1 formulations of bivalent boosters differentially affected the composition of the RBD-specific MBC population (**Fig. 5d-f** and **Extended Data Fig. 10 c,d**). The cross-reactivity of IgGs secreted by in vitro stimulated memory B cells to the Wu, BA.1, BQ.1.1 and XBB.1 RBDs was assessed following Wu/BA.1 bivalent vaccination for uninfected individuals and individuals who had experienced an Omicron BT infection (cohorts vii and viii). Vaccination with the Wu/BA.1 bivalent booster did not elicit Omicron RBD-specific MBCs in either the uninfected or Omicron-BT cohorts. Most RBD-specific antibodies were found to be cross-reactive with BA.1 regardless of infection status (mean frequencies: 76.3% in uninfected and 73.6% in Omicron BT infected subjects), whereas a lower fraction cross-reacted with BQ.1.1 (59.9% and 57.4%) and XBB.1 RBDs (55.9% and 57.4%).

Collectively, these data suggest that two Omicron S exposures are not sufficient to overcome the immunological imprinting induced by repeated Wu S exposures but does continue to enrich for MBCs cross-reacting with multiple RBD variants.

## Discussion

SARS-CoV-2 evolution is driven by viral immune escape and transmissibility^60,61^. The Omicron BA.1, BA.2 and BA.5 lineages, which led to successive waves of infection over the past year, were characterized by immune evasion from Wu S-elicited neutralizing antibodies, higher ACE2-binding affinity, impaired S-mediated fusogenicity and a change in entry pathway due to reduced TMPRSS2-dependence^3–15,39^. Here, we report that the recently emerged SARS-CoV-2 Omicron variants have an unprecedented level of immune evasion with reduction of neutralizing antibody titers reaching up to 100-fold for XBB.1. Whereas BQ.1.1 and BA.2.75.2 retained the high ACE2 binding affinity, similar to earlier Omicron variants, XBB.1 has a lower affinity, comparable to that of the Wu RBD. Although XBB.1 is the most immune evasive of these three Omicron variants, its reduced affinity for ACE2 relative to other co-circulating strains may have hindered its spread. The enhanced ACE2 binding affinity of the newly emerging XBB.1.5, which harbors the S486P RBD mutation (relative to BA.2), may explain the current spread of this rapidly rising variant^47^. We also demonstrate that BQ.1.1 and XBB.1 S display an increased resistance to protease inhibitors blocking membrane or endosomal viral entry relative to other Omicron lineages, suggesting peculiar alterations in the viral entry process of these variants. Collectively, these findings illustrate the complex interplay of immune evasion, fusogenicity and ACE2 affinity in driving SARS-CoV-2 evolution.

There is growing evidence that multiple mechanisms can contribute to protection from infection and severe disease, including humoral and cellular responses^26,62–65^. In particular, antibody-dependent cell cytotoxicity (ADCC) and antibody-dependent opsonophagocytosis are Fc-mediated effector functions that can promote virus clearance and enhance adaptive immune responses in vivo, independently of direct viral neutralization. In this context, it is intriguing that both neutralizing and binding antibody titers were reported as correlates of protection in a phase 3 study^66^. We recently showed that sotrovimab (S309) retained in vitro effector functions against BA.2 and conferred Fc-dependent protection in the lungs of mice infected with BA.2^67^. This preclinical finding is supported by the accumulation of real-world evidence suggesting that adverse clinical outcomes, such as hospitalization and mortality, remained consistently low during the BA.2 wave in patients treated with sotrovimab^68–72^. Here, we show that sotrovimab retained in vitro effector functions against all Omicron variants at levels comparable to those observed with Wu. Importantly, despite 94-fold reduced in vitro neutralizing activity, prophylactic administration of S309 (sotrovimab parent) protected mice against BQ.1.1 challenge, suggesting that Fc-dependent effector functions can compensate for substantial losses of neutralizing activity. Our observation that vaccine-elicited polyclonal plasma antibodies cross-react and trigger Fc*γ*RIIIa signaling, as a surrogate of ADCC, upon recognition of the BQ.1.1 and XBB.1 S glycoproteins, concur with observations made with prior Omicron variants^73–75^ and further hint at a protective role for broadly reactive antibodies with effector functions. Our findings are consistent with a model in which the erosion of neutralizing antibody titers due to viral immune evasion is associated with increased frequency of breakthrough infections, while the persistence of functional cross-reactive antibodies contributes to protection against severe COVID-19^76^.

Immune imprinting, also termed original antigenic sin, was first described in 1960 based on the observation that influenza virus infection with a strain distinct from prior exposures preferentially boosted antibody responses against epitopes shared with the original strain instead of inducing responses against the new strain^77^. Analysis of the specificity of antibodies secreted by plasmablasts obtained approximately 1-2 weeks following infection with the antigenically-shifted 2009 H1N1 swine origin pandemic influenza virus revealed a recall of pre-existing, cross-reactive memory B cells^78,79^. Conversely, antibodies secreted by plasma cells and memory B cells^80^ obtained from the same individuals after seasonal influenza vaccination, corresponding to a second antigenic exposure, were subtype-specific (i.e., targeting non-conserved epitopes). We and others previously showed that Omicron breakthrough infections of Wu-vaccinated subjects primarily recall cross-reactive memory B cells specific for epitopes shared by multiple SARS-CoV-2 variants rather than priming naïve B cells recognizing Omicron RBD-specific epitopes^1,43,51^. Unexpectedly, we observed a low to undetectable number of MBCs specific for Omicron RBDs, and not cross-reacting with the Wu RBD, even after two exposures to Omicron S antigens, including with Wu/BA.5 or Wu/BA.1 bivalent mRNA vaccination. This observation may be explained by a combination of strong immune imprinting and by the antigenic dominance of Wu antigens in bivalent vaccines. This mechanism is reminiscent of current efforts to develop universal influenza vaccines to elicit cross-reactive antibodies by serial stimulation with heterologous chimeric hemagglutinins^81^. As expected, however, we found that bivalent Wu/BA.5 mRNA vaccination enriches for MBCs that are cross-reactive with the vaccine-matched and mismatched RBD variants, relative to monovalent Wu mRNA vaccination. The identification of suitable strategies to design broadly protective vaccines against sarbecoviruses^21,22,82–88^ and the development of mucosal vaccines^89–91^ may result in the development of variant-proof SARS-CoV-2 vaccines through the elicitation of functional cross-reactive antibodies.

## Methods

### Cells and Viruses

Cell lines used in this study were obtained from ATCC (HEK293T and VeroE6), Thermo Fisher Scientific (Expi-CHO-S cells, FreeStyle 293-F cells and Expi293F cells), Takara (Lenti-X 293T cells), kindly gifted from Jesse Bloom (HEK293T-ACE2), or generated in lab (BHK-21-GFP_1-10_, VeroE6-TMPRSS2 or VeroE6-TMPRSS2-GFP_11_). None of the cell lines used were authenticated nor tested for mycoplasma contamination. SARS-CoV-2 isolates used in this study were obtained through BEI Resources, NIAID, NIH: (Wild type isolate hCoV-19/USA-WA1/2020, NR-52281 deposited by the Centers for Disease Control and Prevention ; Lineage B.1.1.529, BA.2; Omicron Variant Isolate hCoV-19/USA/CO-CDPHE-2102544747/2021, NR-56520; Lineage XBB.1.5; Omicron Variant Isolate hCoV-19/USA/MD-HP40900/2022, NR-59104, contributed by Dr. Andrew S. Pekosz). Viruses were propagated and titered on Vero-TMPRSS2 cells in house. The genomic sequences of all strains were confirmed by Sanger and/or NGS sequencing.

### Sample donors

Samples were obtained from SARS-CoV-2 convalescent and vaccinated individuals under study protocols approved by the local institutional review boards (Canton Ticino and Canton Aargau Ethics Committees, Switzerland). All donors provided written informed consent for the use of blood and blood derivatives (such as peripheral blood mononuclear cells, sera or plasma) for research. Sera and peripheral blood mononuclear cells (PBMCs) from individuals who received the Wuhan-Hu-1/BA.5 bivalent mRNA vaccines were obtained from the HAARVI study approved by the University of Washington Human Subjects Division Institutional Review Board (STUDY00000959). Demographic data for these individuals is presented in **Supplementary Tables 5 and 6**.

### Constructs

The full-length Wuhan-Hu-1/G614, Delta, BA.1, BA.2, and BA.4/5 S constructs with a 21 amino acid C-terminal deletion used for pseudovirus were previously described elsewhere (). The full-length BA.2.75.2 and XBB.1 S constructs containing a 21 amino acid C-terminal deletion were codon-optimized, synthesized, and inserted the HDM vector by Genscript. The full-length BQ.1.1 S construct containing a 21 amino acid C-terminal deletion was generated by mutagenesis of the BA.4/5 S construct by Genscript.

The SARS-CoV-2 Wuhan-Hu-1 RBD construct containing an N-terminal mu-phosphatase secretion signal and a C-terminal octa-histidine tag followed by flexible linker and Avi tag was previously described elsewhere. The BA.4/5 RBD construct containing an N-terminal BM40 secretion tag and a C-terminal octa-histidine tag followed by flexible linker and Avi tag was previously described elsewhere. The BA.2.75.2, BQ.1.1, and BA.4/5 RBD constructs containing an N-terminal BM40 secretion tag and a C-terminal octa-histidine tag followed by flexible linker and Avi tag were codon optimized, synthesized, and inserted into the pcDNA3.1(+) vector by Genscript. The boundaries of the construct are N-328RFPN331 and 528KKST531-C.

### Generation of VSV pseudovirus

Replication defective VSV pseudovirus expressing SARS-CoV-2 spike proteins corresponding to the ancestral Wuhan-Hu-1 virus and the VOCs were generated as previously described (*3*), with some modifications. Lenti-X 293T cells (Takara) were seeded in 15-cm^2^ dishes at a density of 10 × 10^6^ cells per dish and the following day were transfected with 25 µg of spike expression plasmid with TransIT-Lenti (Mirus, 6600) according to the manufacturer’s instructions. One day after transfection, cells were infected with VSV-luc (VSV-G) with a multiplicity of infection (MOI) of 3 for 1 h, rinsed three times with PBS containing Ca2+ and Mg2+, then incubated for an additional 24 h in complete medium at 37 °C. The cell supernatant was clarified by centrifugation, aliquoted, and frozen at -80 °C. Spike expression plasmids used for the generation of VSV pseudoviruses carry the following mutations: BA.1: A67V, Δ69-70, T95I, G142D, Δ143-145, Δ211, L212I, ins214EPE, G339D, S371L, S373P, S375F, K417N, N440K, G446S, S477N, T478K, E484A, Q493R, G496S, Q498R, N501Y, Y505H, T547K, D614G, H655Y, N679K, P681H, N764K, D796Y, N856K, Q954H, N969K, L981F; <u>BA.2</u>: T19I, L24-, P25-, P26-, A27S, G142D, V213G, G339D, S371L, S373P, S375F, D405N, R408S, K417N, N440K, S477N, T478K, E484A, Q493R, Q498R, N501Y, Y505H, D614G, H655Y, N679K, P681H, N764K, D796Y, N856K, Q954H, N969K; K417N, N440K, G446S, N460K, S477N, T478K, E484A, Q498R, N501Y, Y505H, D614G, H655Y, N679K, P681H, N764K, D796Y, Q954H, N969K; <u>BA.2.75.2</u>: T19I, L24-, P25-, P26-, A27S, G142D, K147E, W152R, F157L, I210V, V213G, G257S, G339H, R346T, S371F, S373P, S375F, T376A, D405N, R408S, K417N, N440K, G446S, N460K, S477N, T478K, E484A, F486S, Q498R, N501Y, Y505H, D614G, H655Y, N679K, P681H, N764K, D796Y, Q954H, N969K, D1199N; <u>BQ.1</u>: T19I, L24-, P25-, P26-, A27S, Δ69-70, G142D, V213G, G339D, S371F, S373P, S375F, T376A, D405N, R408S, K417N, N440K, K444T, L452R, N460K, S477N, T478K, E484A, F486V, Q498R, N501Y, Y505H, D614G, H655Y, N679K, P681H, N764K, D796Y, Q954H, N969K; <u>BQ.1.1</u>: T19I, L24-, P25-, P26-, A27S, Δ69-70, G142D, V213G, G339D, R436T, S371F, S373P, S375F, T376A, D405N, R408S, K417N, N440K, K444T, L452R, N460K, S477N, T478K, E484A, F486V, Q498R, N501Y, Y505H, D614G, H655Y, N679K, P681H, N764K, D796Y, Q954H, N969K; <u>BF.7</u>: T19I, L24-, P25-, P26-, A27S, Δ69-70, G142D, V213G, G339D, R436T, S371F, S373P, S375F, T376A, D405N, R408S, K417N, N440K, L452R, S477N, T478K, E484A, F486V, Q498R, N501Y, Y505H, D614G, H655Y, N679K, P681H, N764K, D796Y, Q954H, N969K; <u>XBB.1</u>: T19I, L24-, P25-, P26-, A27S, V83A, G142D, Y144-, H146Q, Q183E, V213E, G252V, G339H, R346T, L368I, S371F, S373P, S375F, T376A, D405N, R408S, K417N, N440K, V445P, G446S, N460K, S477N, T478K, E484A, F486S, F490S, Q498R, N501Y, Y505H, D614G, H655Y, N679K, P681H, N764K, D796Y, Q954H, N969K; <u>XBB.1.5</u>: T19I, L24-, P25-, P26-, A27S, V83A, G142D, Y144-, H146Q, Q183E, V213E, G252V, G339H, R346T, L368I, S371F, S373P, S375F, T376A, D405N, R408S, K417N, N440K, V445P, G446S, N460K, S477N, T478K, E484A, F486P, F490S, Q498R, N501Y, Y505H, D614G, H655Y, N679K, P681H, N764K, D796Y, Q954H, N969K; CH.1.1: T19I, del24-26, A27S, G142D, K147E, W152R, F157L, I210V, V213G, G257S, G339H, R346T, S371F, S373P, S375F, T376A, D405N, R408S, K417N, N440K, K444T, G446S, L452R, N460K, S477N, T478K, E484A, F486S, Q498R, N501Y, Y505H, D614G, H655Y, N679K, P681H, N764K, D796Y, Q954H, N969K Pseudotyped VSV was produced as previously described. Briefly, HEK293T were split into poly-D-lysine coated 15 cm plates and grown overnight until they reached approximately 70-80% confluency. The cells were washed 3 times with Opti-MEM (Gibco) and transfected with either the Wuhan-Hu-1(D614), Wu-G614, Delta, BA.1, BA.2, BA.4/5, BA.2.75.2, BQ.1.1, or XBB.1 S constructs using Lipofectamine 2000 (Life Technologies). After 4-6 hours, the media was supplemented with an equal volume of DMEM supplemented with 20% FBS and 2% PS. The cells were incubated for 20-24 hours, washed 3 times with DMEM, and infected with VSVΔG-luc. Two hours after VSVΔG-luc infection, the cells were then washed an additional five times with DMEM. The cells were grown in DMEM (for virus stocks used in sotrovimab neutralization assays) or in DMEM supplemented with anti-VSV-G antibody (I1-mouse hybridoma supernatant diluted 1:25, from CRL-2700, ATCC) for 18-24 hours, after which the supernatant was harvested and clarified by low-speed centrifugation at 2,500 g for 10 min. The supernatant was then filtered (0.45 μm) and some virus stocks were concentrated 10 times using a 30 kDa centrifugal concentrator (Amicon Ultra). The pseudotyped viruses were then aliquoted and frozen at -80°C.

### VSV pseudovirus neutralization

Vero E6 cells were grown in DMEM supplemented with 10% FBS and seeded into white-walled 96 well plates (PerkinElmer, 6005688) at a density of 20,000 cells per well. The next day, monoclonal antibodies were serially diluted in pre-warmed complete medium, mixed with pseudoviruses and incubated for 1 h at 37 °C in round bottom polypropylene plates. Medium from cells was aspirated and 50 µl of virus–monoclonal antibody complexes were added to cells, which were then incubated for 1 h at 37 °C. An additional 100 µl of pre-warmed complete medium was then added on top of complexes and cells were incubated for an additional 16–24 h. Conditions were tested in duplicate or triplicate wells on each plate and 6-8 wells per plate contained untreated infected cells (defining the 0% of neutralization, ‘MAX RLU’ value) and uninfected cells (defining the 100% of neutralization, ‘MIN RLU’ value). Virus–monoclonal antibody-containing medium was then aspirated from cells and 50 or 100 µl of a 1:2 dilution of SteadyLite Plus (PerkinElmer) or BioGlo (Promega) in PBS with Ca^2+^ and Mg^2+^ was added to cells. Plates were incubated for 15 min at room temperature and then analyzed on the Synergy-H1 (Biotek). The average relative light units (RLUs) of untreated infected wells (MAX RLUave) were subtracted by the average of MIN RLU (MIN RLUave) and used to normalize percentage of neutralization of individual RLU values of experimental data according to the following formula: (1 - (RLUx – MIN RLUave)/ (MAX RLUave – MIN RLUave)) × 100. Data were analyzed with Prism (v.9.1.0). IC_50_ values were calculated from the interpolated value from the log(inhibitor) versus response, using variable slope (four parameters) nonlinear regression with an upper constraint of <100. Each neutralization experiment was conducted on at least two independent experiments – that is, biological replicates – in which each biological replicate contains a technical duplicate or triplicate. VeroE6-TMPRSS2 were split into white-walled, clear bottom 96 well plates (Corning) and grown overnight until they reached approximately 70% confluency. Sera (or plasma) were diluted in DMEM starting at a 1:33 or 1:66 dilution and serially diluted in DMEM at a 1:3 dilution thereafter. Pseudotyped VSV was diluted at a 1:25 to 1:100 ratio in DMEM and an equal volume was added to the diluted sera. The virus-sera mixture was incubated for 30 minutes at room temperature and added to the VeroE6-TMPRSS2 cells. After two hours, an equal volume of DMEM supplemented with 20% FBS and 2% PS was added to the cells. After 20-24 hours, ONE-Glo EX (Promega) was added to each well and the cells were incubated for 5 minutes at 37°C. Luminescence values were measured using a BioTek plate reader. Luminescence readings from the neutralization assays were normalized and analyzed using GraphPad Prism 9. The relative light unit (RLU) values recorded from uninfected cells were used to define 100% neutralization and RLU values recorded from cells infected with pseudovirus without sera or antibodies were used to define 0% neutralization. ID50 were determined from the normalized data points using a [inhibitor] vs. normalized response – variable slope model using at least two technical repeats to generate the curve fits. At least two biological replicates with two distinct batches of pseudovirus were conducted for each sample.

### Neutralization of authentic SARS-CoV-2 viruses

Vero-TMPRSS2 cells were seeded into black-walled, clear-bottom 96-well plates at 20,000 cells/well and cultured overnight at 37°C. The next day, 9-point 4-fold serial dilutions of mAbs were prepared in growth media (DMEM + 10% FBS). The different SARS-CoV-2 strains were diluted in infection media (DMEM + 2% BSA) at a final MOI of 0.01 PFU/cell, added to the mAb dilutions and incubated for 30 min at 37°C. Media was removed from the cells, mAb-virus complexes were added and incubated at 37°C for 18h (wild-type, XBB.1.5) or 24h (BA.2). Cells were fixed with 4% PFA (Electron Microscopy Sciences, #15714S), permeabilized with Triton X-100 (SIGMA, #X100-500ML) and stained with an antibody against the viral nucleocapsid protein (Sino Biologicals, #40143-R001) followed by a staining with the nuclear dye Hoechst 33342 (Fisher Scientific, # H1399) and a goat anti-rabbit Alexa Fluor 647 antibody (Invitrogen, #A-21245). Plates were imaged on a Cytation5 plate reader. Whole well images were acquired (12 images at 4X magnification per well) and nucleocapsid-positive cells were counted using the manufacturer’s software.

### Pseudotyped VSV entry assays with protease inhibitors

VeroE6-TMPRSS2 or HEK293T-ACE2 were split into white-walled, clear bottom 96 well plates (Corning) at a density of 18,000 or 36,000 cells, respectively, and grown overnight. The following day, the growth media was removed and, for assays conducted with VeroE6-TMPRSS2, the cells washed once with DMEM. The cells were incubated for 2 hours with DMEM containing 50 µM of Camostat (Sigma), Nafamostat (Sigma), E-64d (Sigma), or 0.5% DMSO. All three protease inhibitors were dissolved in DMSO to a concentration of 10 mM and diluted in DMEM. The protease inhibitors were removed and pseudovirus diluted 1:50 or 1:200 in DMEM was added to the cells. After two hours, an equal volume of DMEM supplemented with 20% FBS and 2% PS was added to the cells. After 20-24 hours, ONE-Glo EX (Promega) was added to each well and the cells were incubated for 5 minutes at 37°C. Luminescence values were measured using a BioTek plate reader. Luminescence readings from the neutralization assays were normalized and analyzed using GraphPad Prism 9. The RLU values recorded from uninfected cells were used to define 0% infectivity and RLU values recorded from cells incubated with 0.5% DMSO only and infected with pseudovirus were used to define 100% infectivity. Twelve technical replicates were performed for each inhibitor and pseudovirus and at least two biological replicates with two distinct batches of pseudovirus were conducted.

### Recombinant protein production for BLI, FACS, and cryoEM

SARS-CoV-2 RBD proteins were produced and purified from Expi293F cells as previously described. In brief, cells were grown to a density of 3 × 10^6^ cells/mL and transfected using the ExpiFectamine 293 Transfection Kit (ThermoFisher Scientific). Three to 5 days post-transfection, proteins were purified from clarified supernatants using HisTrap HP affinity columns (Cytiva) and washed with ten column volumes of 20 mM imidazole, 25 mM sodium phosphate pH 8.0, and 300 mM NaCl before elution on a gradient to 500 mM imidazole, 25 mM sodium phosphate pH 8.0, and 300 mM NaCl. Proteins were buffer exchanged into 20 mM sodium phosphate pH 8 and 100 mM NaCl and concentrated using centrifugal filters (Amicon Ultra) before being flash frozen. For cryo-EM, recombinant ACE2 (residues 19-615 from Uniprot Q9BYF1 with a C-terminal thrombin cleavage site-TwinStrep-10xHis-GGG-tag, and N-terminal signal peptide) was expressed in ExpiCHO-S cells at 37°C and 8% CO_2_ with kifunensine added to 10 µM. Cell culture supernatant was collected eight days post transfection, supplemented with buffer to a final concentration of 80 mM Tris-HCl pH 8.0, 100 mM NaCl, and then incubated with BioLock (IBA GmbH) solution. ACE2 was purified using a 5 mL StrepTrap HP column (Cytiva) followed by isolation of the monomeric ACE2 by size exclusion chromatography using a Superdex 200 Increase 10/300 GL column (Cytiva) pre-equilibrated in PBS. Recombinant S309 Fab used for cryo-EM was expressed in HEK293 suspension cells, purified using CaptureSelect IgG-CH1 resin and buffer exchanged into PBS (ATUM Bio).

### Biolayer interferometry

Biotinylated Wu, BA.4/5, BA.2.75.2, BQ.1.1, and XBB.1 RBDs were diluted to a concentration of 5 ng/µL in 10X kinetics buffer and loaded onto pre-hydrated streptavidin biosensors to a 1 nm total shift. The loaded tips were dipped into a 1:3 dilution series of monomeric human ACE2 starting at 900 nM or 300 nM for 300 seconds followed by dissociation in 10X kinetics buffer for 300 seconds. All steps of the affinity measurements using biolayer interferometry were carried out at 30°C with a shaking speed of 1,000 rpm.

### Surface plasmon resonance to measure binding of RBDs with ACE2

Measurements were performed using a Biacore T200 instrument. Experiments were performed at 25°C, with the samples held at 15°C in the instrument prior to injection. A CM5 chip with covalently immobilized anti-Avi polyclonal antibody (GenScript, Cat #: A00674-40) was used for surface capture of His-Avi tag containing RBDs. Running buffer was 1x HBS-EP+ pH 7.4 (10 mM HEPES, 150 mM NaCl, 3 mM EDTA and 0.05% v/v Surfactant P20) (Cytiva, Cat #: BR100669). Experiments were performed with a 4-point dilution series of monomeric S309 Fab or monomeric Strep-tagged ACE2 starting at 300nM. The dilution factors are 3-fold (300, 100, 33.3, 11,1nM), 3.25-fold (300, 92.3, 28.4, 8.7nM) or 4-fold (300, 75, 18.8, 4.7nM). Experiments were run as single-cycle kinetics, n=2-14 for each RBD ligand. Data were double reference-subtracted and fit to a binding model using Biacore Evaluation software. The 1:1 binding model was used to determine the kinetic parameters. Kinetic values out of the instrument’s limit were omitted. K_D_ values are reported as the average of all replicates with the corresponding standard deviation (**Supplementary Table 2**).

### In vivo studies

Animal studies were carried out in accordance with the recommendations in the Guide for the Care and Use of Laboratory Animals of the National Institutes of Health. The protocols were approved by the Institutional Animal Care and Use Committee at the Washington University School of Medicine (assurance number A3381–01). Virus inoculations were performed under anesthesia that was induced and maintained with ketamine hydrochloride and xylazine, and all efforts were made to minimize animal suffering. Heterozygous K18-hACE2 C57BL/6 J mice (strain: 2B6.Cg-Tg(K18-ACE2)2Prlmn/J) were obtained from The Jackson Laboratory. All animals were housed in groups of 3 to 5 and fed standard chow diets. The photoperiod was 12 h on:12 h off dark/light cycle. The ambient animal room temperature was 70° F, controlled within ±2° and the room humidity was 50%, controlled within ±5%. Eight- to ten-week-old female K18-hACE2 mice were administered indicated doses of S309 or control (anti-WNV hE16) mAb dose by intraperitoneal injection one day before or after intranasal inoculation with 10^4^ FFU of BQ.1.1. Weight was recorded daily, and animals were euthanized on day +6 after virus inoculation.

### Measurement of viral RNA levels

Tissues were weighed and homogenized with zirconia beads in a MagNA Lyser instrument (Roche Life Science) in 1 mL of DMEM medium supplemented with 2% heat-inactivated FBS. Tissue homogenates were clarified by centrifugation at approximately 10,000 × *g* for 5 min and stored at -80 °C. RNA was extracted using the MagMax mirVana Total RNA isolation kit (Thermo Fisher Scientific) on the Kingfisher Flex extraction robot (Thermo Fisher Scientific). RNA was reverse transcribed and amplified using the TaqMan RNA-to-CT 1-Step Kit (Thermo Fisher Scientific). Reverse transcription was carried out at 48 °C for 15 min followed by 2 min at 95 °C. Amplification was accomplished over 50 cycles as follows: 95 °C for 15 s and 60 °C for 1 min. Copies of SARS-CoV-2 *N* gene RNA in samples were determined using a previously published assay<u>^42^</u>. Briefly, a TaqMan assay was designed to target a highly conserved region of the *N* gene (Forward primer: ATGCTGCAATCGTGCTACAA; Reverse primer: GACTGCCGCCTCTGCTC; Probe: /56-FAM/TCAAGGAAC/ZEN/AACATTGCCAA/3IABkFQ/). This region was included in an RNA standard to allow for copy number determination down to 10 copies per reaction. The reaction mixture contained final concentrations of primers and probe of 500 and 100 nM, respectively.

### Viral plaque assay

VeroE6-TMPRSS2-ACE2 cells were seeded at a density of 1 × 10^5^ cells per well in 24-well tissue culture plates. The following day, medium was removed and replaced with 200 μL of material to be titrated diluted serially in DMEM supplemented with 2% FBS. One hour later, 1 mL of methylcellulose overlay was added. Plates were incubated for 72 h, and then fixed with 4% paraformaldehyde (final concentration) in PBS for 20 min. Plates were stained with 0.05% (w/v) crystal violet in 20% methanol and washed twice with distilled, deionized water.

### Transient expression of recombinant SARS-CoV-2 protein and flow cytometry

ExpiCHO-S cells were seeded at 6 × 10^6^ cells/mL in a volume of 5 mL in a 50 mL bioreactor. The following day, cells were transfected with SARS-CoV-2 spike glycoprotein-encoding pcDNA3.1(+) plasmids (BetaCoV/Wuhan-Hu-1/2019, accession number MN908947, Wu-D614; Omicron BA.2, BQ.1.1, XBB.1 and BA.2-E340A generated by overlap PCR mutagenesis of the Wuhan D614 plasmid) harboring the Δ19 C-terminal truncation. Spike encoding plasmids were diluted in cold OptiPRO SFM (Life Technologies, 12309-050), mixed with ExpiFectamine CHO Reagent (Life Technologies, A29130) and added to cells. Transfected cells were then incubated at 37°C with 8% CO2 with an orbital shaking speed of 250 RPM (orbital diameter of 25 mm) for 24 to 48 h. Transiently transfected ExpiCHO-S cells were harvested and washed twice in wash buffer (PBS 2% FBS, 2mM EDTA). Cells were counted and distributed into round bottom 96-well plates (Corning, 3799) and incubated with serial dilutions of mAb starting at 10 μg/mL. Alexa Fluor647-labelled Goat anti-human IgG secondary Ab (Jackson ImmunoResearch, 109–606–098) was prepared at 2 μg/mL and added onto cells after two washing steps. Cells were then washed twice and resuspended in wash buffer for data acquisition at Ze5 cytometer (BioRad).

### Fc-mediated effector functions (ADCC)

Primary cells were collected from healthy human donors with informed consent and authorization via the Comitato Etico Canton Ticino (Switzerland). ADCC assays were performed using ExpiCHO-S cells transiently transfected with SARS-CoV-2 spike glycoproteins (Wuhan D614, BA.2, BQ.1.1 or XBB.1) as targets. NK cells were isolated from fresh blood of healthy donors using the MACSxpress NK Isolation Kit (Miltenyi Biotec, cat. no. 130-098-185). Target cells were incubated with titrated concentrations of mAbs for 10 min and then with primary human NK cells at an effector to target ratio ranging from 7.75:1 to 9:1. ADCC was measured using the LDH release assay (Cytotoxicity Detection Kit (LDH) (Roche; cat. no. 11644793001) after 4 h incubation at 37°C.

### Measurement of plasma effector functions

Antibody-dependent activation of human FcγRIIIa by plasma antibodies was performed with a bioluminescent reporter assay. ExpiCHO-S cells transiently expressing full-length SARS-CoV-2 S from Wuhan-D614, BA.5, BQ.1.1 and XBB.1 (target cells) were incubated with serial dilutions of plasma from immune donors. After a 20-minute incubation, Jurkat reporter cells stably expressing FcγRIIIa V158 and NFAT-driven luciferase gene (effector cells) were added at an effector to target ratio of 6:1 for FcγRIIIa. Signaling was quantified by the luciferase signal produced via activation of the NFAT pathway. Luminescence was measured after 22 hours of incubation at 37°C with 5% CO_2_ with a luminometer using the Bio-Glo-TM Luciferase Assay Reagent according to the manufacturer’s instructions (Promega).

### Antigen-specific memory B cell repertoire analysis (AMBRA) of secreted IgGs

Replicate cultures of total unfractionated PBMCs obtained from SARS-CoV-2 infected and/or vaccinated individuals were seeded in 96 U-bottom plates (Corning) in RPMI1640 supplemented with 10% fetal calf serum (Hyclone), sodium pyruvate, MEM non-essential amino acids, stable glutamine, 2-mercaptoethanol, Penicillin-Streptomycin, Kanamycin and Transferrin. Memory B cell stimulation and differentiation was induced by adding 2.5 μg/ml R848 (3 M) and 1000 U/ml human recombinant IL-2 at 37 °C and 5% CO_2_. After 10 days, the cell culture supernatants were collected for ELISA analysis.

### Enzyme-linked immunosorbent assay (ELISA)

Ninety-six half area well-plates (Corning, 3690) were coated overnight at 4 °C with 25 μl of sarbecovirus RBD proteins prepared at 5 μg/ml in PBS pH 7.2. Plates were then blocked with PBS 1% BSA (Sigma-Aldrich, A3059) and subsequently incubated with mAb serial dilutions for 1 h at room temperature. After 4 washing steps with PBS 0.05% Tween 20 (PBS-T) (Sigma-Aldrich, 93773), goat anti-human IgG secondary antibody (Southern Biotech, 2040-04) was added and incubated for 1 h at room temperature. Plates were then washed four times with PBS-T and 4-nitrophenyl phosphate (pNPP, Sigma-Aldrich, 71768) substrate was added. After 30 min incubation, absorbance at 405 nm was measured by a plate reader (Biotek) and data were plotted using Prism GraphPad 9.1.0. To test MBC-derived antibodies, Spectraplate-384 with high protein binding treatment (custom made from Perkin Elmer) were coated overnight at 4°C with 3 µg/ml of RBD (produced in house), SARS-CoV RBD (produced in house), Omicron BA.1, BQ.1.1 and XBB:1 RBDs (produced in house) in PBS pH 7.2 or PBS alone as control. Plates were subsequently blocked with Blocker Casein (1%) in PBS (Thermo Fisher Scientific, 37528) supplemented with 0.05% Tween 20 (Sigma Aldrich, 93773-1KG). The coated plates were incubated with diluted B cell supernatant for 1h at RT. Plates were washed with PBS containing 0.05 % Tween20 (PBS-T), and binding was revealed using secondary goat anti-human IgG-AP (Southern Biotech, 2040-04). After washing, pNPP substrate (Sigma-Aldrich, 71768-25G) was added and plates were read at 405 nm after 1 h or 30 minutes.

### Blockade of RBD binding to human ACE2

Memory B cell culture supernatants were diluted in PBS and mixed with SARS-CoV-2 RBD mouse Fc-tagged antigen (Sino Biological, 40592-V05H, final concentration 20 ng/ml) and incubated for 30 min at 37°C. The mix was added for 30 min to ELISA 384-well plates (NUNC, P6366-1CS) pre-coated overnight at 4°C with 4 µg/ml human ACE2 (produced in house) in PBS. Plates were washed with PBS containing 0.05 % Tween20 (PBS-T), and RBD binding was revealed using secondary goat anti-mouse IgG-AP (Southern Biotech, 1032-04). After washing, pNPP substrate (Sigma-Aldrich, 71768-25G) was added and plates were read at 405 nm after 1h.

### Recombinant protein production for SPR binding assays and AMBRA ELISA

SARS-CoV-2 RBD plasmids encode for residues 328-531 of the spike protein from GenBank NC_045512.2 with an N-terminal signal peptide and a C-terminal 8xHis-AviTag or thrombin cleavage site-8xHis-AviTag. Proteins were expressed in Expi293F cells (Thermo Fisher Scientific) at 37 °C and 8% CO_2_. Transfections were performed using the ExpiFectamine 293 Transfection Kit (Thermo Fisher Scientific). Cell culture supernatants were collected four to five days after transfection and supplemented with 10x PBS to a final concentration of 2.5x PBS (342.5 mM NaCl, 6.75 mM KCl and 29.75 mM phosphates). SARS-CoV-2 RBDs were purified by IMAC using Cobalt or Nickel resin followed by buffer exchange into PBS using Amicon centrifugal filters (Milipore Sigma) or by size exclusion chromatography using a Superdex 200 Increase 10/300 GL column (Cytiva). For SPR binding measurements, recombinant ACE2 (residues 19-615 from Uniprot Q9BYF1 with a C-terminal thrombin cleavage site-TwinStrep-10xHis-GGG-tag, and N-terminal signal peptide) was expressed in Expi293F cells at 37°C and 8% CO_2_. Transfection was performed using the ExpiFectamine 293 Transfection Kit (Thermo Fisher Scientific). Cell culture supernatant was collected seven days after transfection, supplemented to a final concentration of 80 mM Tris-HCl pH 8.0, 100 mM NaCl, and then incubated with BioLock solution (IBA GmbH). ACE2 was purified using a 1 mL StrepTrap HP column (Cytiva) followed by isolation of the monomeric ACE2 by size exclusion chromatography using a Superdex 200 Increase 10/300 GL column (Cytiva) pre-equilibrated in PBS. Recombinant S309 Fab used for SPR binding studies was produced in ExpiCHO-S cells and purified using a Capture Select CH1-XL MiniChrom Column (ThermoFisher), followed by buffer exchange into PBS using a HiPrep 26/10 Desalting Column (Cytiva).

### Surface plasmon resonance (SPR) assays to measure binding of the S309 Fab to RBDs

Measurements were performed using a Biacore T200 instrument. A CM5 chip with covalently immobilized StrepTactin XT was used for surface capture of Twin-Strep Tag-containing RBDs. Running buffer was HBS-EP+ pH 7.4 (Cytiva) and measurements were performed at 25°C. Experiments were performed with a 3-fold dilution series of monomeric S309 Fab at 300, 100, 33 and 11 nM and were run as single-cycle kinetics. Data were double reference-subtracted and fit to a binding model using Biacore Evaluation software. The 1:1 binding model was used to estimate the kinetics parameters. The experiment was performed twice with two biological replicates for each ligand (RBDs). K_D_ values are reported as the average of two replicates with the corresponding standard deviation.

### Cell-cell fusion assay

Cell-cell fusion assays using a split GFP system was conducted as previously described. In brief, VeroE6-TMPRSS2-GFP_11_ cells were split into 96-well, glass bottom, black walled plates (CellVis) at a density of 36,000 cells per well. BHK-21-GFP_1-10_ cells were split into 6-well plates at a density of 1 × 10^6^ cells per well. The following day, the growth media was removed and replaced with DMEM containing 10% FBS and 1% PS and the cells were transfected with 4 µg of S protein using Lipofectamine 2000. Twenty-four hours after transfection, BHK-21-GFP_1-10_ expressing the S protein were washed three times using FluoroBrite DMEM (Thermo Fisher) and detached using an enzyme-free cell dissociation buffer (Gibco). The VeroE6-TMPRSS2-GFP_11_ were washed three times with FluoroBrite DMEM and 9,000 BHK-21-GFP_1-10_ cells were plated on top of the VeroE6-TMPRSS2-GFP_11_ cells. The cells were incubated at 37°C and 5% CO_2_ in a Cytation 7 plate Imager (Biotek) and both brightfield and GFP images were collected every 30 minutes for 18 hours. Fusogenicity was assessed by measuring the area showing GFP fluorescence for each image using Gen5 Image Prime v3.11 software.

To measure surface expression of the variant SARS-CoV-2 S protein, 1 × 10^6^ transiently transfected BHK-21-GFP_1-10_ cells were collected by centrifugation at 1,000 x g for 5 min. The cells were washed once with PBS and fixed with 2% paraformaldehyde. The cells were washed twice with flow staining buffer (1% BSA, 1 mM EDTA, 0.1% NaN_3_ in PBS) and labeled with 250 µg/mL of S2L20, an NTD-directed antibody that recognizes all currently and previously circulating SARS-CoV-2 variants, for 45 minutes. The cells were washed three times with flow staining buffer and labeled with a PE-conjugated anti-Human IgG Fc antibody (Thermo Fisher) for 30 mins. The cells were washed an additional three times and resuspended in flow staining buffer. The labeled cells were analyzed using a BD FACSymphony A3. ells were gated on singleton events and a total of 10,000 singleton events were collected for each sample. The fraction of S-positive cells was determined in FlowJo 10.8.1 by gating singleton events for the mock transfected cells on PE intensity.

### Flow cytometry analysis of SARS-CoV-2 RBD-reactive MBCs

RBD-streptavidin tetramers conjugated to fluorophores were generated by incubating biotinylated Wu, BA.1, BA.2, BA.4/5, or BQ.1.1 with streptavidin at a 4:1 molar ratio for 30 mins at 4*°*C. Excess free biotin was then added to the reaction to bind any unconjugated sites in the streptavidin tetramers. The RBD-streptavidin tetramers were washed once with PBS and concentrated with a 30 kDa centrifugal concentrator (Amicon). An additional streptavidin tetramer conjugated to biotin only was generated and included in the staining.

Approximately 5 to 15 million PMBCs were collected 5-72 days post-vaccination for individuals who received either the Wu monovalent mRNA booster or Wu/BA.5 bivalent mRNA booster. The cells were collected by centrifugation at 1,000 x g for 5 mins at 4*°*C and washed twice with PBS. The cells were then stained with Zombie Aqua dye (Biolegend; diluted 1:100 in PBS) for 30 mins at room temperature after which the cells were washed twice with FACS staining buffer (0.1% BSA, 0.1% NaN_3_ in PBS). The cells were then stained with antibodies for CD20-PECy7 (BD), CD3-Alexa eFluor780 (ThermoFisher), CD8-Alexa eFluor780 (ThermoFisher), CD14-Alexa eFluor780 (ThermoFisher), CD16-Alexa eFluor780 (ThermoFisher), IgM-Alexa Fluor647 (BioLegend), IgD-Alexa Fluor647 (BioLegend), and CD38-Brilliant Violet 785 (BioLegend), all diluted 1:200 in Brilliant Stain Buffer (BD), along with the RBD-streptavidin tetramers for 30 mins at 4*°*C. The cells were washed three times, resuspended in FACS staining buffer, and passed through a 35 µm filter. The cells were examined using a BD FACSAria III and FACSDiva for acquisition and FlowJo 10.8.1 for analysis. Single live CD20^+^/CD3^-^/CD8^-^/CD14^-^/CD16^-^ /IgM^lo^/IgD^lo^/CD38^lo^/RBD^+^ cells were sorted based on reactivity to the Omicron and Wu RBDs into RNAlater and stored at -80*°*C.

### CryoEM sample preparation, data collection and data processing

CryoEM grids of BQ.1.1 RBD-ACE2-S309 or XBB.1 RBD-ACE2-S309 complex were prepared fresh after SEC purification. For BQ.1.1 RBD-ACE2-S309 complex, three microliters of 0.25 mg/ml BQ.1.1 RBD-ACE2-S309 were loaded onto freshly glow discharged R 2/2 UltrAuFoil grids, prior to plunge freezing using a vitrobot MarkIV (ThermoFisher Scientific) with a blot force of 0 and 6 sec blot time at 100 % humidity and 22°C. Data were acquired using an FEI Titan Krios transmission electron microscope operated at 300 kV and equipped with a Gatan K3 direct detector and Gatan Quantum GIF energy filter, operated in zero-loss mode with a slit width of 20 eV. For BQ.1.1 RBD-ACE2-S309 data set, automated data collection was carried out using Leginon at a nominal magnification of 105,000x with a pixel size of 0.843 Å and stage tilt angle of 30°. 6,487 micrographs were collected with a defocus range comprised between -0.5 and -2.5 μm. For XBB.1 RBD-ACE2-S309 complex, samples were prepared using the Vitrobot Mark IV (Thermo Fisher Scientific) with R 2/2 UltrAuFoil grids and with a Chameleon (SPT Labtech) with self-wicking nanowire Cu R1.2/0.8 holey carbon grids. For XBB.1 RBD-ACE2-S309 data set, 6,355 micrographs from UltrAuFoil grids and 2,889 micrographs from chameleon grids were collected with a defocus range comprised between -0.2 and -3 μm. The dose rate was adjusted to 15 counts/pixel/s, and each movie was acquired in super-resolution mode fractionated in 75 frames of 40 ms. Movie frame alignment, estimation of the microscope contrast-transfer function parameters, particle picking, and extraction were carried out using Warp.

Two rounds of reference-free 2D classification were performed using cryoSPARC to select well-defined particle images. These selected particles were subjected to two rounds of 3D classification with 50 iterations each (angular sampling 7.5° for 25 iterations and 1.8° with local search for 25 iterations) using Relion with an initial model generated with ab-initio reconstruction in cryoSPARC. 3D refinements were carried out using non-uniform refinement along with per-particle defocus refinement in CryoSPARC. Selected particle images were subjected to the Bayesian polishing procedure implemented in Relion3.0 before performing another round of non-uniform refinement in cryoSPARC followed by per-particle defocus refinement and again non-uniform refinement. To further improve the density of the BQ.1.1 RBD and XBB.1 RBD, the particles were subjected to focus 3D classification without refining angles and shifts using a soft mask encompassing the ACE2, RBD and S309 variable domains using a tau value of 60 in Relion. Particles belonging to classes with the best resolved local density were selected and subjected to non-uniform refinement using cryoSPARC. Local resolution estimation, filtering, and sharpening were carried out using CryoSPARC. Reported resolutions are based on the gold-standard Fourier shell correlation (FSC) of 0.143 criterion and Fourier shell correlation curves were corrected for the effects of soft masking by high-resolution noise substitution.

### Model building and refinement

UCSF Chimera and Coot were used to fit atomic models into the cryoEM maps. Spike RBD domain, ACE2, S309 Fab models were refined and relaxed using Rosetta using sharpened and unsharpened maps.

### Statistical Analysis

All statistical tests were performed as described in the indicated figure legends using Prism v9.0. The number of independent experiments performed are indicated in the relevant figure legends.

## Data Availability

All datasets generated and information presented in the study are available from the corresponding authors on reasonable request. Materials generated in this study can be available on request and may require a material transfer agreement.

## Acknowledgements

We would like to thank Abigail E. Powell, Yvonne M. DaCosta, John M. Errico and Ashley Lin for assistance with protein production. This study was supported by the National Institute of Allergy and Infectious Diseases (DP1AI158186 and HHSN272201700059C to D.V.), the National Institute of Health Cellular and Molecular Biology Training Grant (T32GM007270 to A.A.), a Pew Biomedical Scholars Award (D.V.), an Investigators in the Pathogenesis of Infectious Disease Awards from the Burroughs Wellcome Fund (D.V.), Fast Grants (D.V.), the University of Washington Arnold and Mabel Beckman cryoEM center and the National Institute of Health grant S10OD032290 (to D.V.). D.V. is an Investigator of the Howard Hughes Medical Institute. O.G. is funded by the Swiss Kidney Foundation. We acknowledge the Research Council of Cantonal Hospital Aarau for the financial support (to M.J.K.).

## Author Contributions

A.A, L.P., D.C. and D.V. designed the experiments; A.A, K.S., M.B., H.D., J.B., C.S.F., F.M., D.P., C.Sa., M.G., R.A., D.J., C.M., E.D., C.S, C.Y., A.R., S.Su., J.Z., N.F., D.B. and J.N. performed binding, neutralization assays, biolayer interferometry and surface plasmon resonance binding measurements. L.A.P, G.Sc., F.A.L. and D.V. supervised in vitro neutralization assays. E.C. and K.C. designed and performed mutagenesis for the spike mutant expression plasmids. G.L., G.Le., B.G., L.V. and M.A.S. performed experiments to measure effector functions; A.A., C.S.F., M.B., F.M., J.B. and L.P performed memory B cell repertoire analysis. A.A. carried out membrane fusion and protease inhibition experiments. O.G., A.C., P.F., N.F., M.D., O.G., A.C., P.F., A.F.P., M.B., C.G., S.Z., L.B., M.J.K., J.K.L., N.F., and H.C. contributed to the recruitment of donors and collection of plasma samples. J.B.C., S.S. and B.W. performed mouse experiments and viral burden analyses. Y.J.P. carried out cryoEM specimen preparation, data collection and processing. Y.J.P. and D.V. built and refined the atomic models. C.S. purified recombinant glycoproteins. A.A, L.P., G.S., A.L., M.S.D., D.C. and D.V. analyzed the data and wrote the manuscript with input from all authors; M.S.D, D.C., and D.V. supervised the project.

---

**Extended Data Figure 1.**
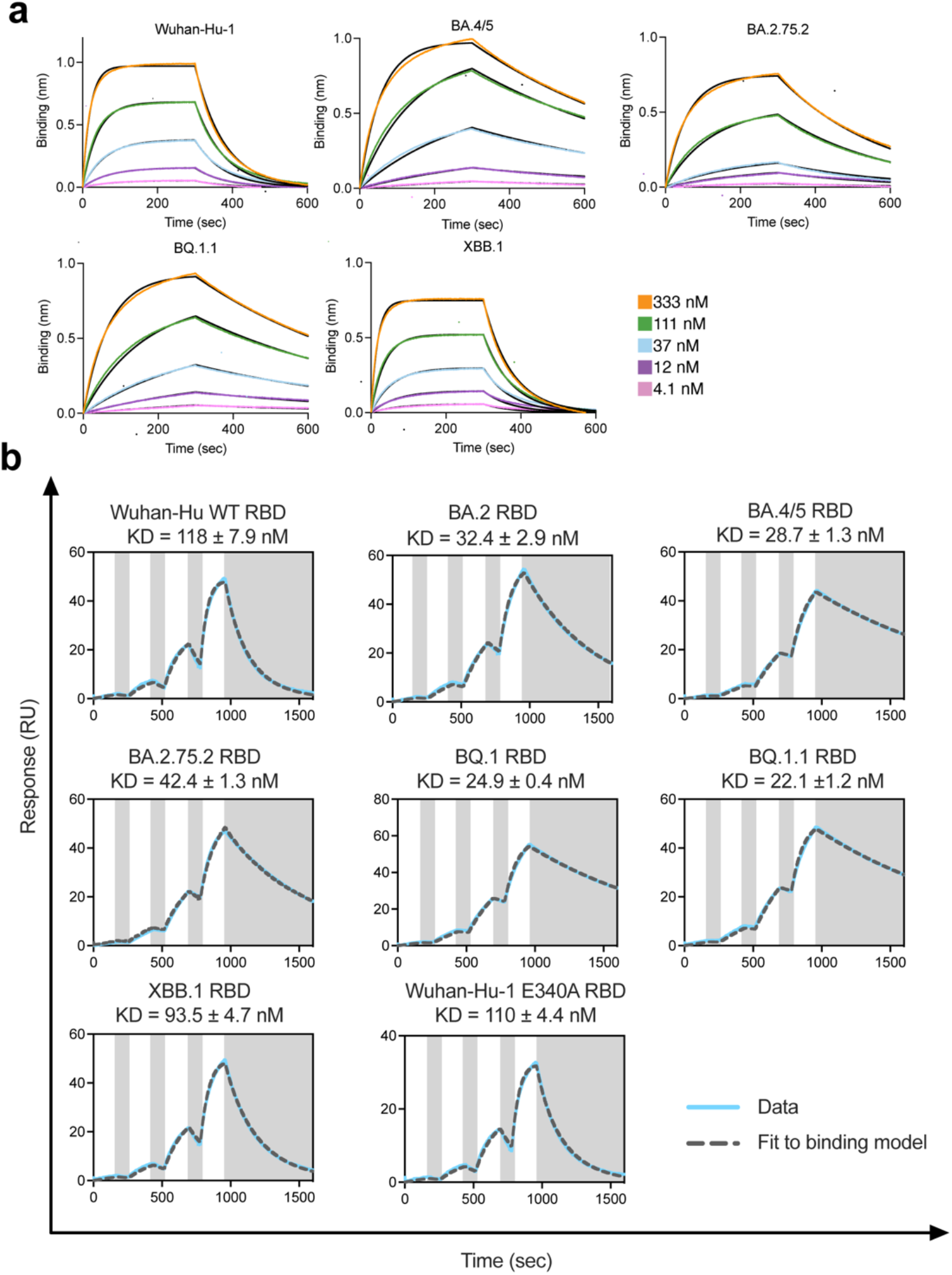
Evaluation of human ACE2 binding to SARS-CoV-2 variant RBDs. **a**, Biolayer interferometry binding curves obtained for monomeric human ACE2 binding to biotinylated Wu, BA.4/5, BA.2.75.2, BQ.1.1, or XBB.1 RBDs immobilized on the surface of streptavidin biosensors. Kinetic rate constants and affinities are shown in **Supplementary Table 1**. Fits are shown as solid black lines. **b**, SPR sensorgrams of monomeric human ACE2 binding to the Wu (Wild-type, WT), BA.2, BA.2.75.2, BA.5, BQ.1, BQ.1.1, XBB.1 and Wu E340A RBDs immobilized at the surface of a SPR chip coated with anti-Avi polyclonal antibody. Experiments were performed with serial dilutions of Fabs and run as single-cycle kinetics. Gray blocks denote the dissociation phase. Fits are shown as dashed grey lines. Kinetic rate constants and affinities are shown in **Supplementary Table 2**.

**Extended Data Figure 2.**
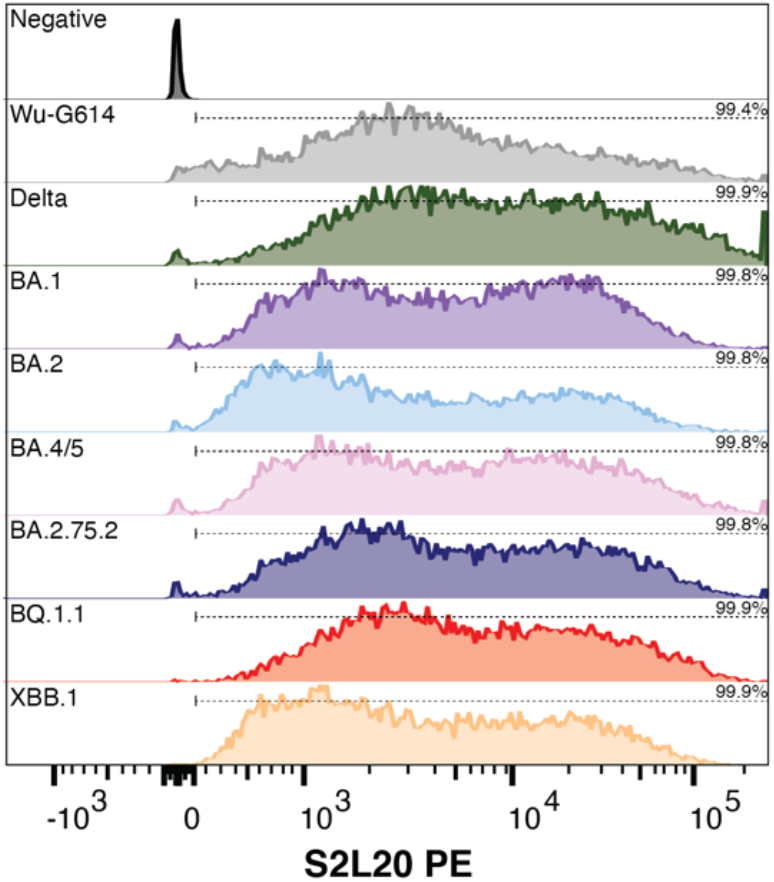
Membrane fusion assay. Quantification of SARS-CoV-2 S surface expression by flow cytometry for BHK-21 GFP_1-10_ cells transfected with Wu-G614, Delta, BA.1, BA.2, BA.4/5, BA.2.75.2, BQ.1.1, or XBB.1 S proteins using the NTD-directed antibody S2L20^24,37^. The y-axis is present as a modal scale scaled to maximum singleton events for that plot. The percentage of S-positive cells is based on the PE intensity for mock transfected (negative) cells and represented by the dashed line above each plot.

**Extended Data Figure 3.**
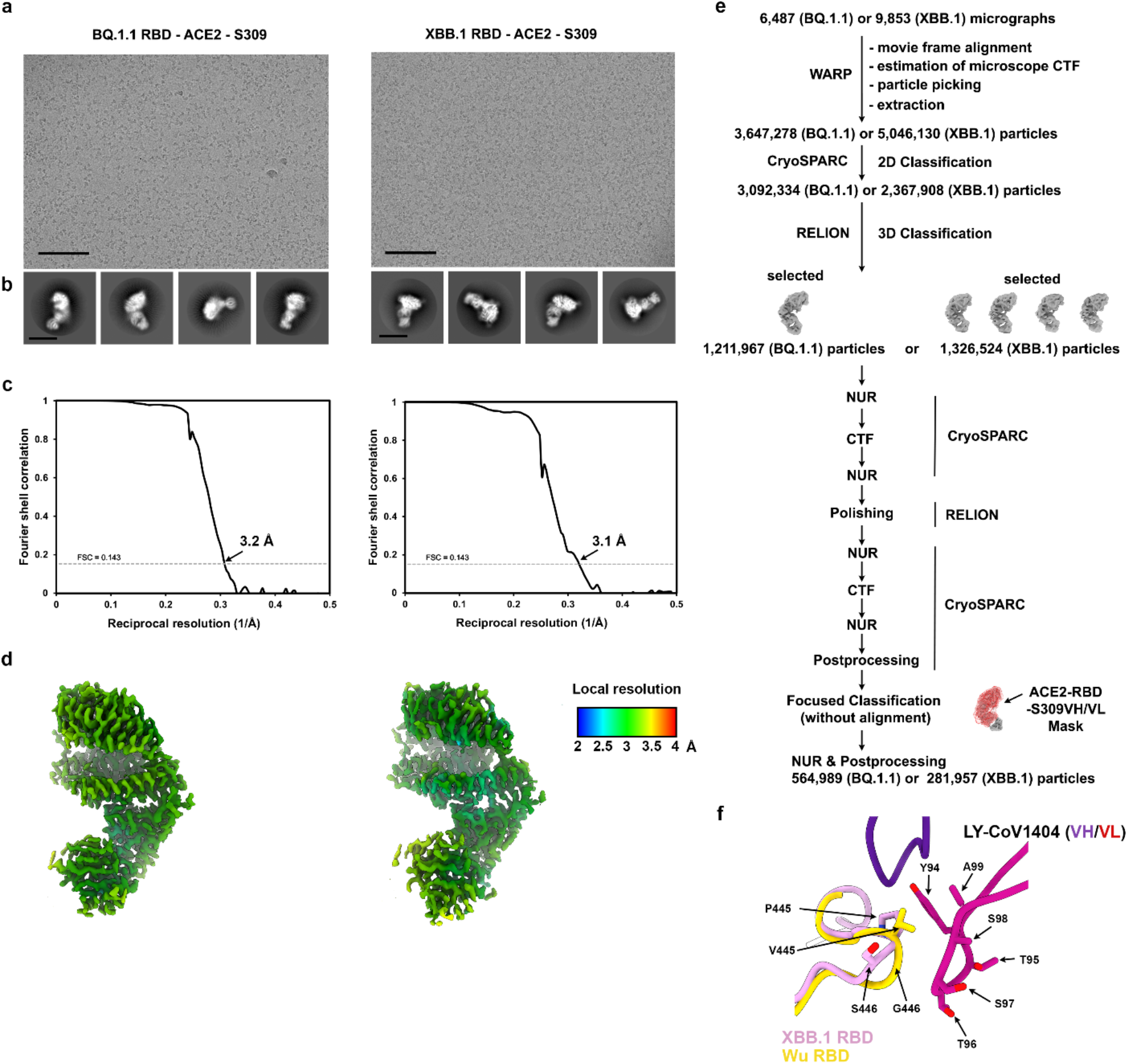
CryoEM data processing of the BQ.1.1 or the XBB.1 RBDs bound to ACE2 and S309. **a-b**, Representative electron micrograph (a) and 2D class averages (b) of the BQ.1.1 RBD (left) or the XBB.1 RBD (right) bound to the human ACE2 ectodomain and the S309 Fab fragment embedded in vitreous ice. The scale bar represents 100nm (a) or 100 Å (b). **c-d**, Gold-standard Fourier shell correlation curves (c) and local resolution maps calculated using cryoSPARC (d) for the 3D reconstructions of the BQ.1.1 RBD (left) or the XBB.1 RBD (right) bound to the human ACE2 ectodomain and the S309 Fab fragment. The 0.143 cutoff is indicated by a horizontal dashed line. **e**, Data processing flowchart. CTF: contrast transfer function; NUR: non-uniform refinement. **f**, Superimposition of the LYCoV1404-bound Wu RBD (gold, PDB 7MMO) crystal structure to the ACE2- and S309-bound XBB.1 RBD (pink) cryoEM structure (S309 and ACE2 are not shown for clarity).

**Extended Data Figure 4.**
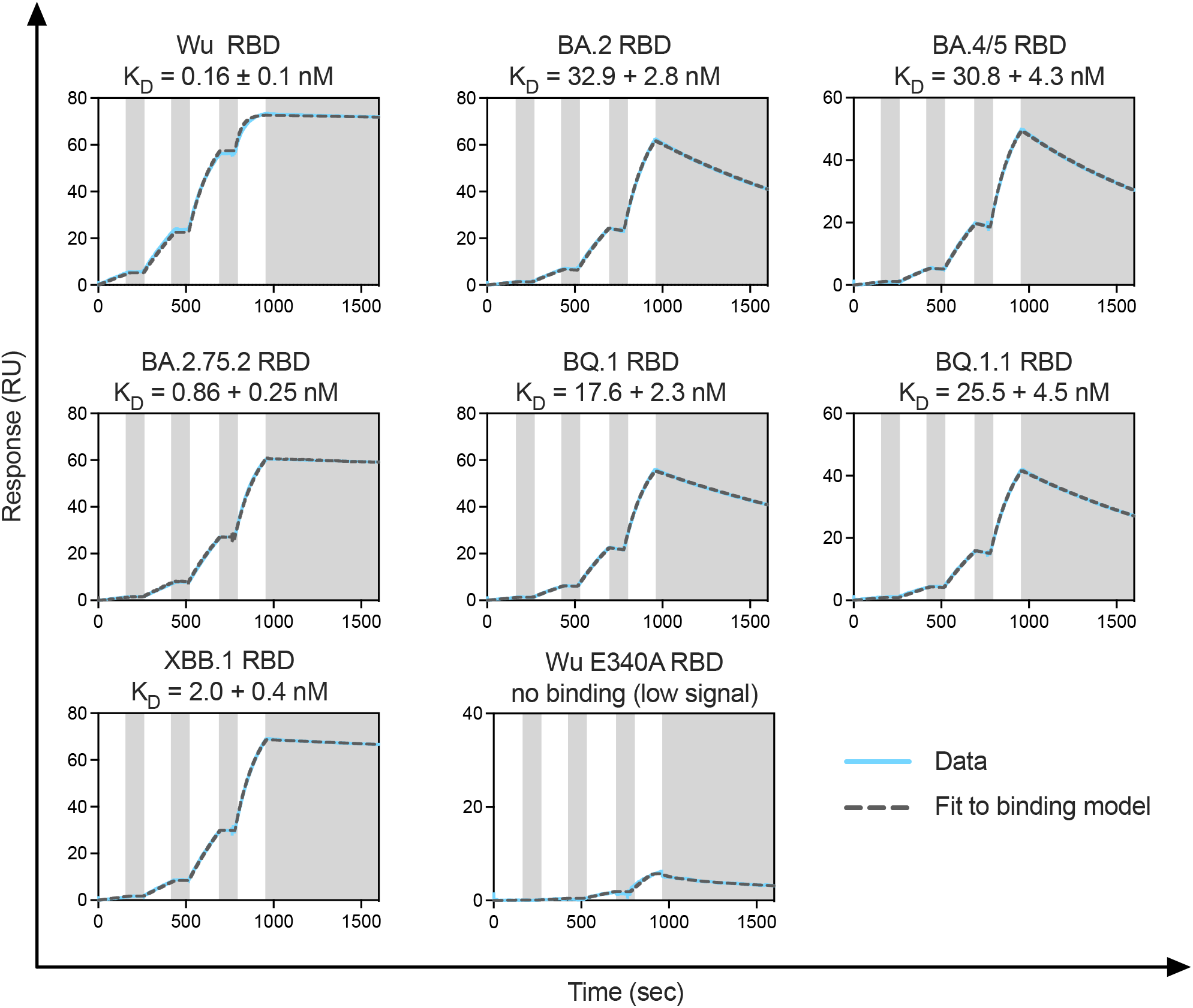
Cross-reactivity of S309 with SARS-CoV-2 variant RBDs. Representative sensograms of S309 Fab binding to the SARS-CoV-2 Wu, BA.2, BA.2.75.2, BA.5, BQ.1, BQ.1.1, XBB.1 and Wu E340A RBDs immobilized at the surface of a SPR chip coated with anti-Avi polyclonal antibody. Experiments were performed with serial dilutions of Fabs and were run as single-cycle kinetics. Gray blocks denote the dissociation phase. Fits are shown as dashed grey lines. Kinetic rate constants and affinities are shown in **Supplementary Table 4**.

**Extended Data Figure 5.**
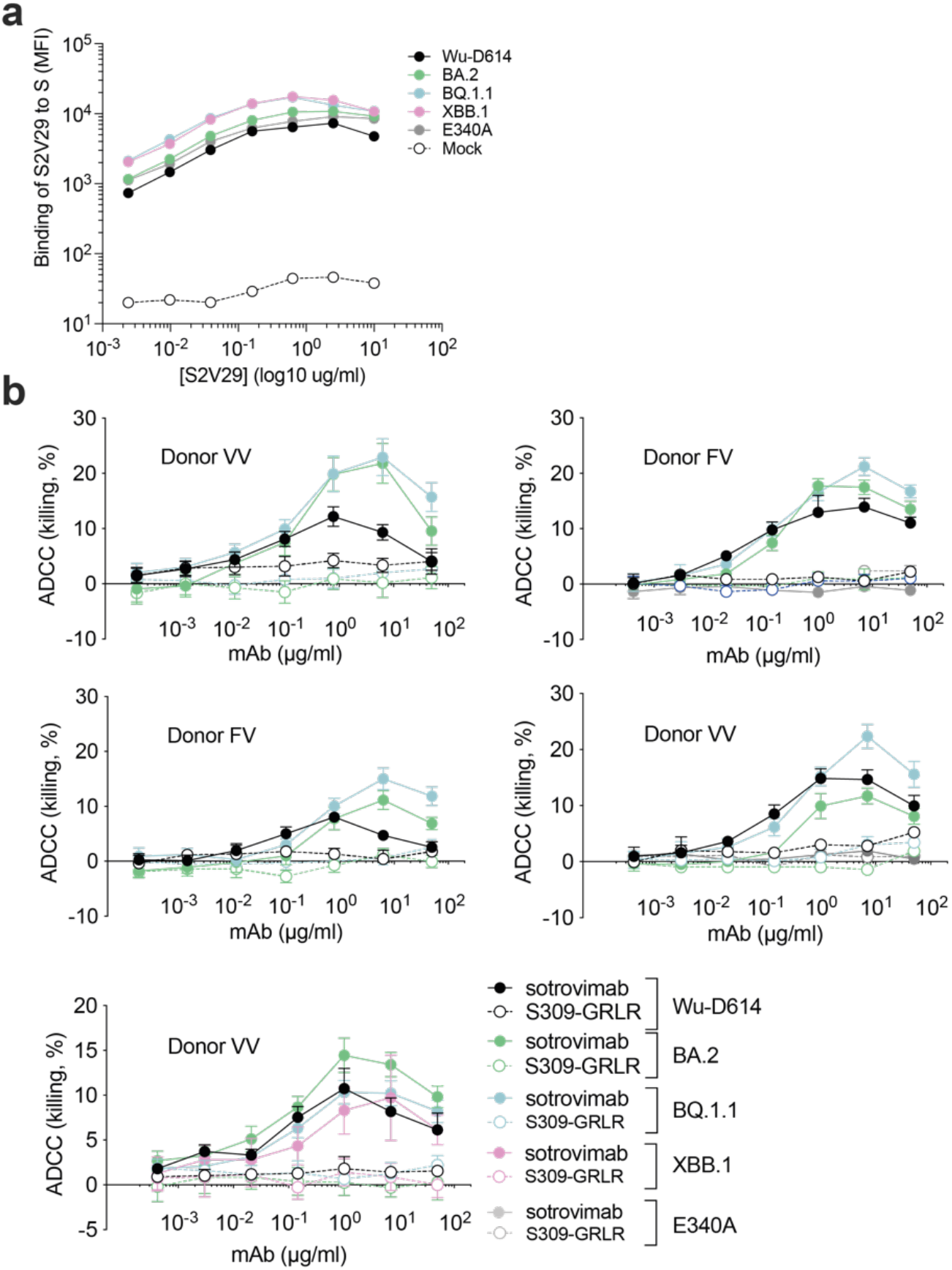
Sotrovimab-mediated ADCC using primary human NK cells. **a**, Binding of the S2V29 monoclonal antibody to SARS-CoV-2 S variants expressed at the surface of ExpiCHO-S cells as measured by flow cytometry. S2V29 is a mAb that retains potent and equal neutralizing activity against Wu-D614, BA.2, BQ.1.1, XBB.1 and E340A VSV pseudoviruses. **b**, ExpiCHO-S cells transiently transfected with expression plasmids encoding Wuhan D614, BA.2, BQ.1.1, XBB.1 and BA.2-E340A S proteins were incubated with the indicated concentrations of sotrovimab or S309-GRLR and mixed with NK cells isolated from healthy donors in a range from 7.75:1 to 9:1 (effector:target). Target cell lysis was determined by a lactate dehydrogenase release assay. Data are presented as mean values +/– standard deviations (SD) from duplicates. Each panel is an individual representative donor. Donors FV, heterozygous for F158 and V158 Fc*γ*RIIIa; VV, homozygous for V158.

**Extended Data Figure 6.**
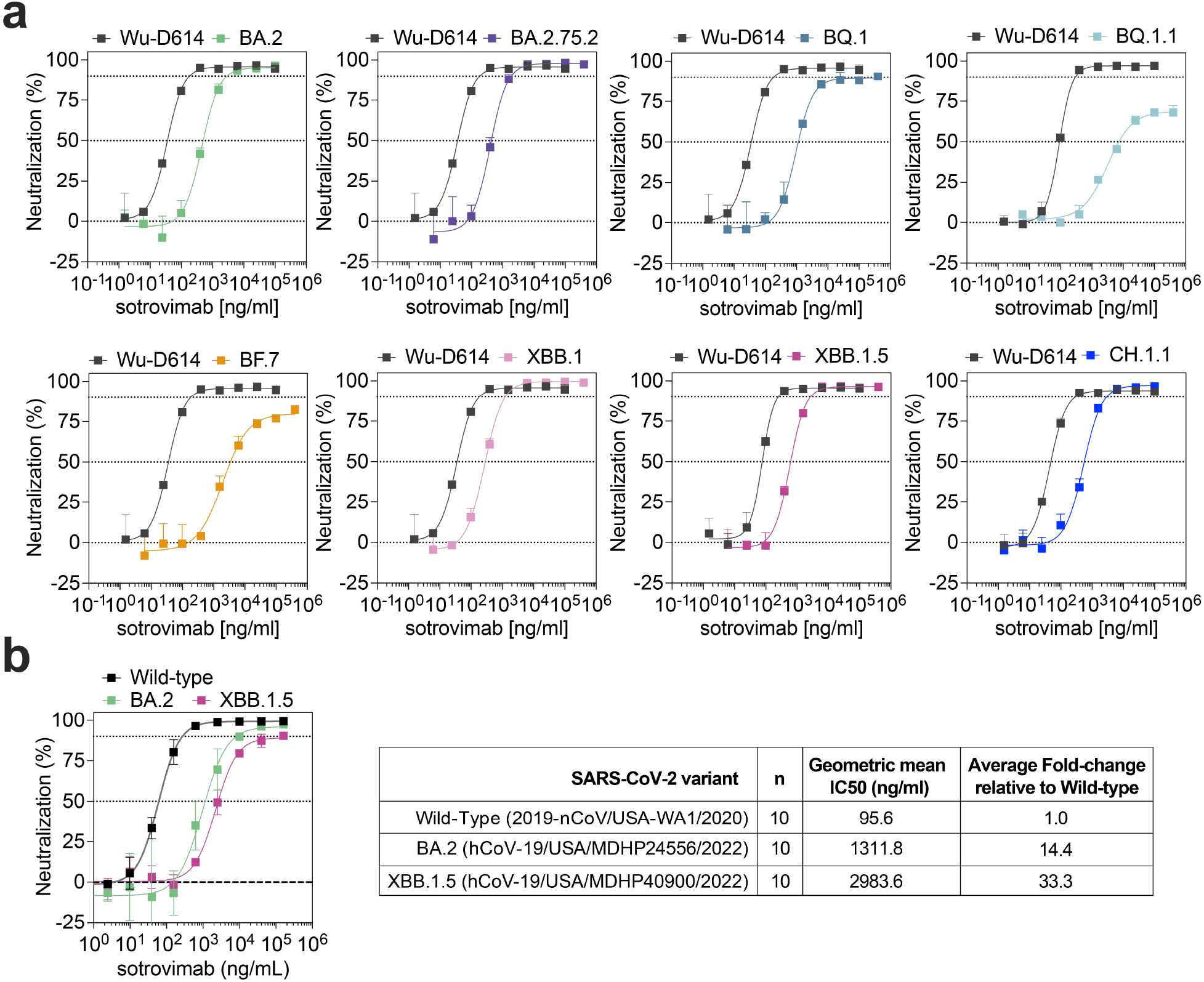
Sotrovimab neutralization of SARS-CoV-2 Omicron variants. **a**, Sotrovimab-mediated neutralization of Wu-D614, BA.2, BA.2.75.2, BQ.1, BQ.1.1, BF.7, XBB.1, XBB.1.5. and CH.1.1 S VSV pseudoviruses using VeroE6 as target cells. Dose-response curves of one representative experiment out of at least 5 experiments are shown. **b**, Sotrovimab-mediated neutralization of Wild-Type (2019-nCoV/USA-WA1/2020), Omicron BA.2 (hCoV-19/USA/MDHP24556/2022) and Omicron XBB.1.5 (hCoV-19/USA/MDHP40900/2022) authentic viruses using VeroE6-TMPRSS2 as target cells. Neutralization data (left panel) represent the means of triplicates ± standard deviation from one representative of n = 10 biologically independent experiments. Shown is also the geometric mean IC_50_ and average fold-change relative to wild-type of the 10 performed experiments (right panel).

**Extended Data Figure 7.**
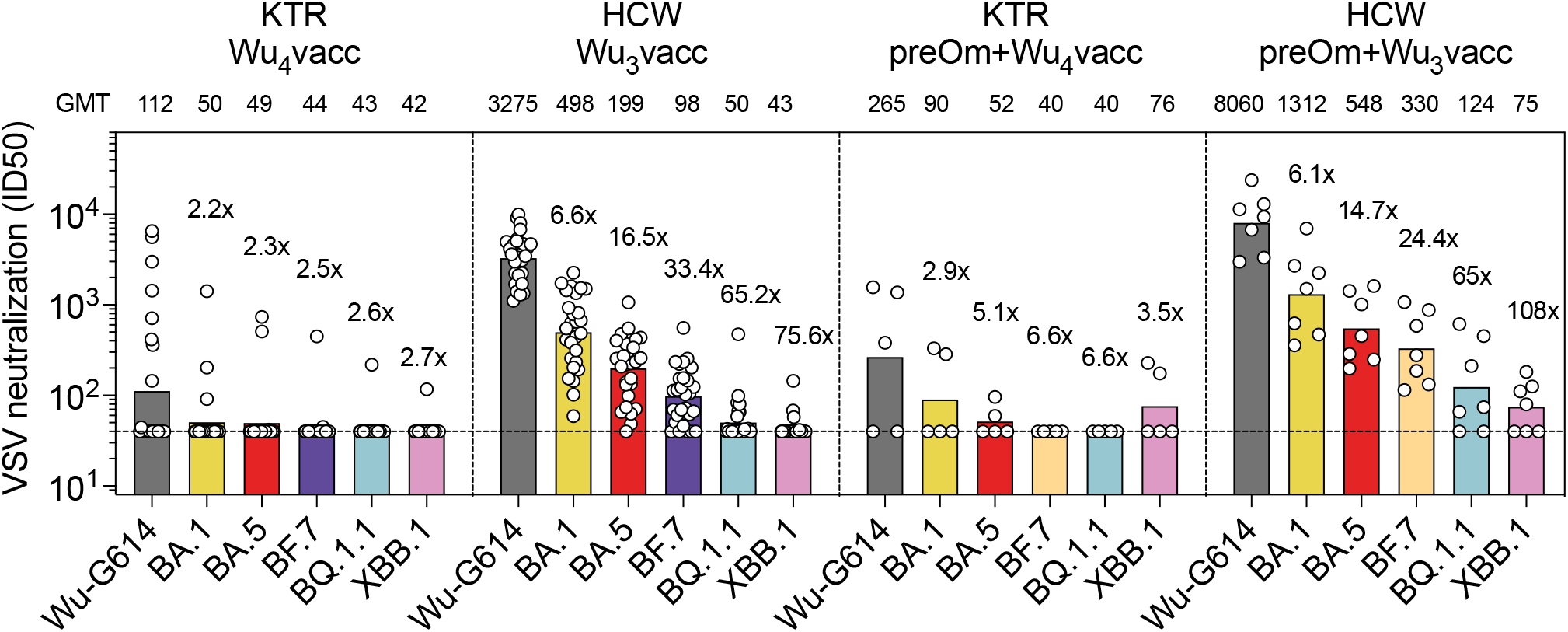
Vaccine-elicited plasma neutralizing antibodies against emerging Omicron variants in kidney transplant recipients. Neutralization of SARS-CoV-2 pseudotyped VSV carrying Wu-G614, Omicron BA.1, BA.5, BF.7, BQ.1.1 and XBB.1 by serum samples from kidney transplant recipients (KTR) or healthcare workers (HCW) collected 2-4 months after receiving 4 (Wu_4_vacc) or 3 doses (Wu_3_vacc) of monovalent Wu vaccine, respectively. Shown are ID50 values from n = 2 technical replicates. Bars and values on top represent geometric mean ID50 titers (GMT). Fold-loss of neutralization against Omicron variants as compared to Wu-G614 is shown above each corresponding bar. Horizontal dashed lines indicate the lowest limit of detectable neutralization in these assays (ID50 = 40). Cohort demographics are summarized in **Supplementary Table 6**.

**Extended Data Figure 8.**
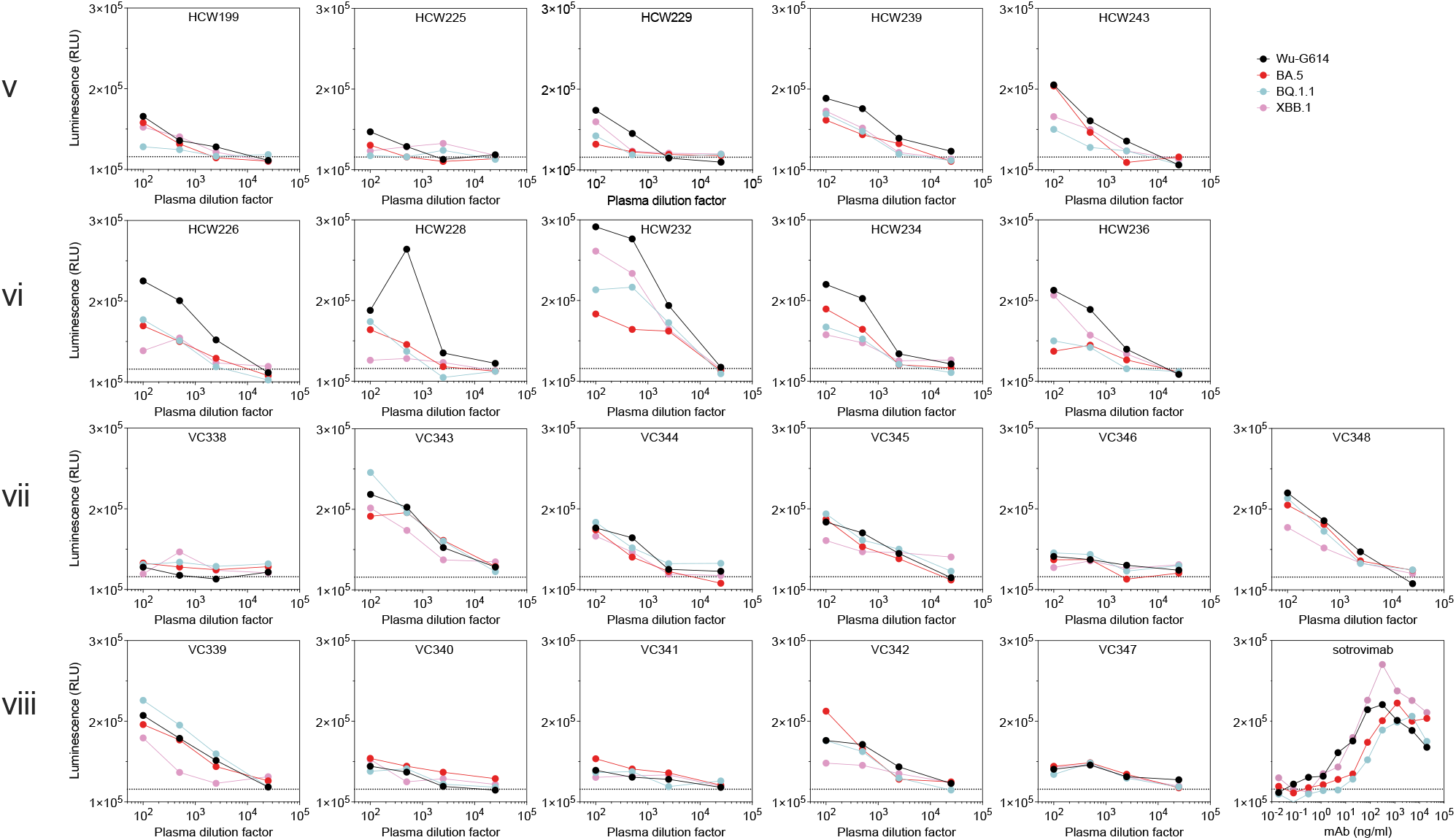
Analysis of individual plasma samples activating Fc*γ*RIIIa. Activation of high-affinity (V158) Fc*γ*RIIIa measured using Jurkat reporter cells and Wu-G614, BA.5, BQ.1.1 and XBB.1 SARS-CoV-2 S glycoprotein-expressing ExpiCHO as target cells. Luminescence (RLU) values from one experiment are shown with plasma samples from cohorts v-viii (n=5 donors for cohort v, n=5 for cohort vi, n=6 for cohort vii and n=5 for cohort viii) and compared to sotrovimab. Horizontal dotted line indicates the lowest limit of detectable activation (RLU = 115,737).

**Extended Data Figure 9.**
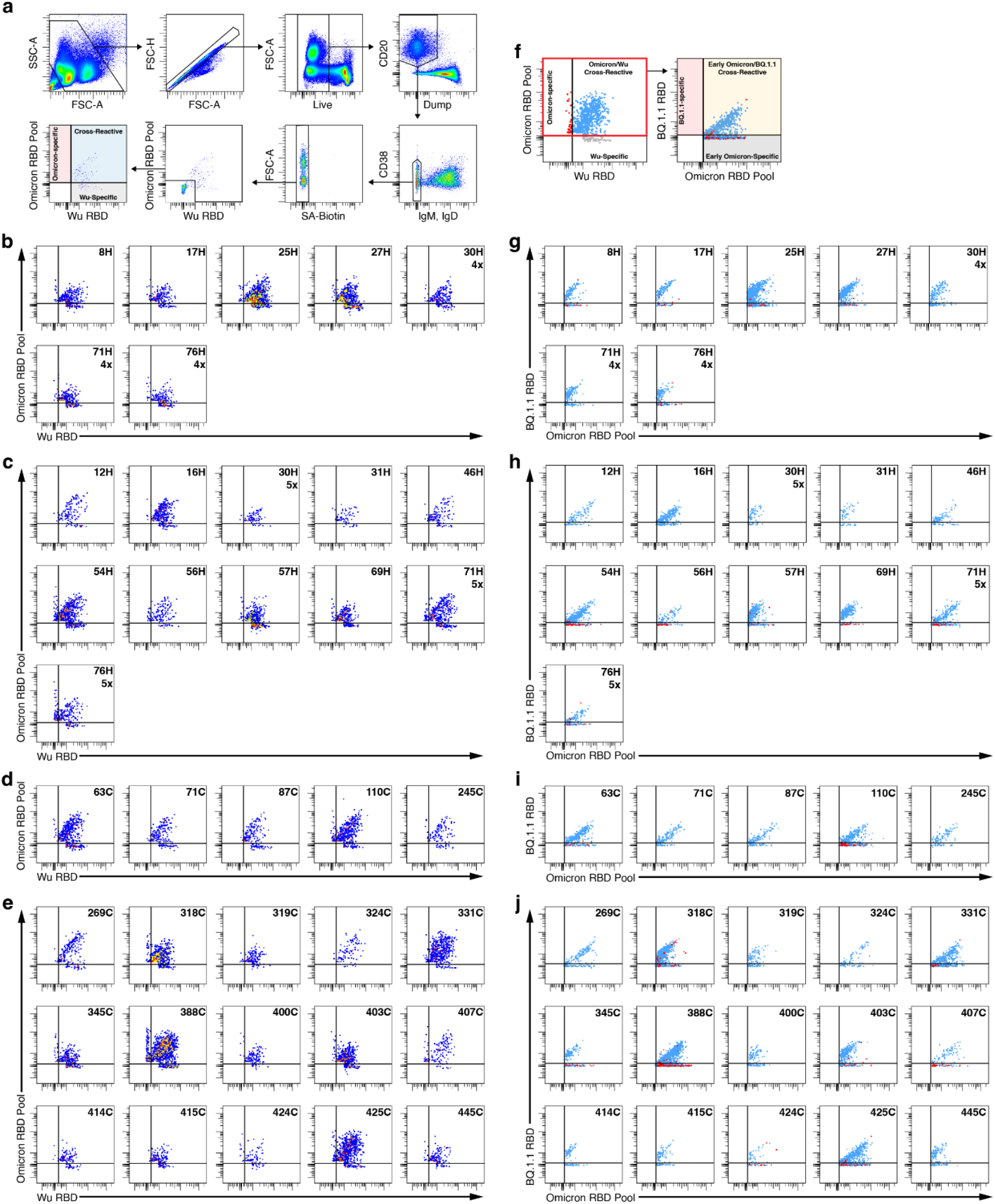
MBC analysis by flow cytometry. **a**, Gating strategy to identify Omicron (BA.1/BA.2/BA.5) pool RBD- and Wu RBD-recognizing MBCs. Dump includes markers for CD3, CD8, CD14, and CD16. Gating for RBD-positive memory B cells was based on staining of PBMCs from healthy donors collected in 2019 prior to the SARS-CoV-2 pandemic. Individual plots showing Omicron (BA.1/BA.2/BA.5) pool RBD- and Wu RBD-positive MBCs for Wu_4_ vaccinated (b), Wu/BA.5 bivalent vaccinated (c), pre-Omicron infected-Wu/BA.5 bivalent vaccinated (d), and Omicron BT-Wu/BA.5 bivalent vaccinated individuals (e). **f**, Gating strategy to determine whether (BA.1, BA.2, and BA.4/5) pool RBD-positive MBCs recognize the BQ.1.1 RBD. Individual plots showing Omicron (BA.1, BA.2, and BA.4/5) RBD pool and BQ.1.1 RBD-recognizing memory B cells for Wu4 vaccinated (g), Wu/BA.5 bivalent vaccinated (h), pre-Omicron infected-Wu/BA.5 bivalent vaccinated (i), and Omicron BT-Wu/BA.5 bivalent vaccinated individuals (j).

**Extended Data Figure 10.**
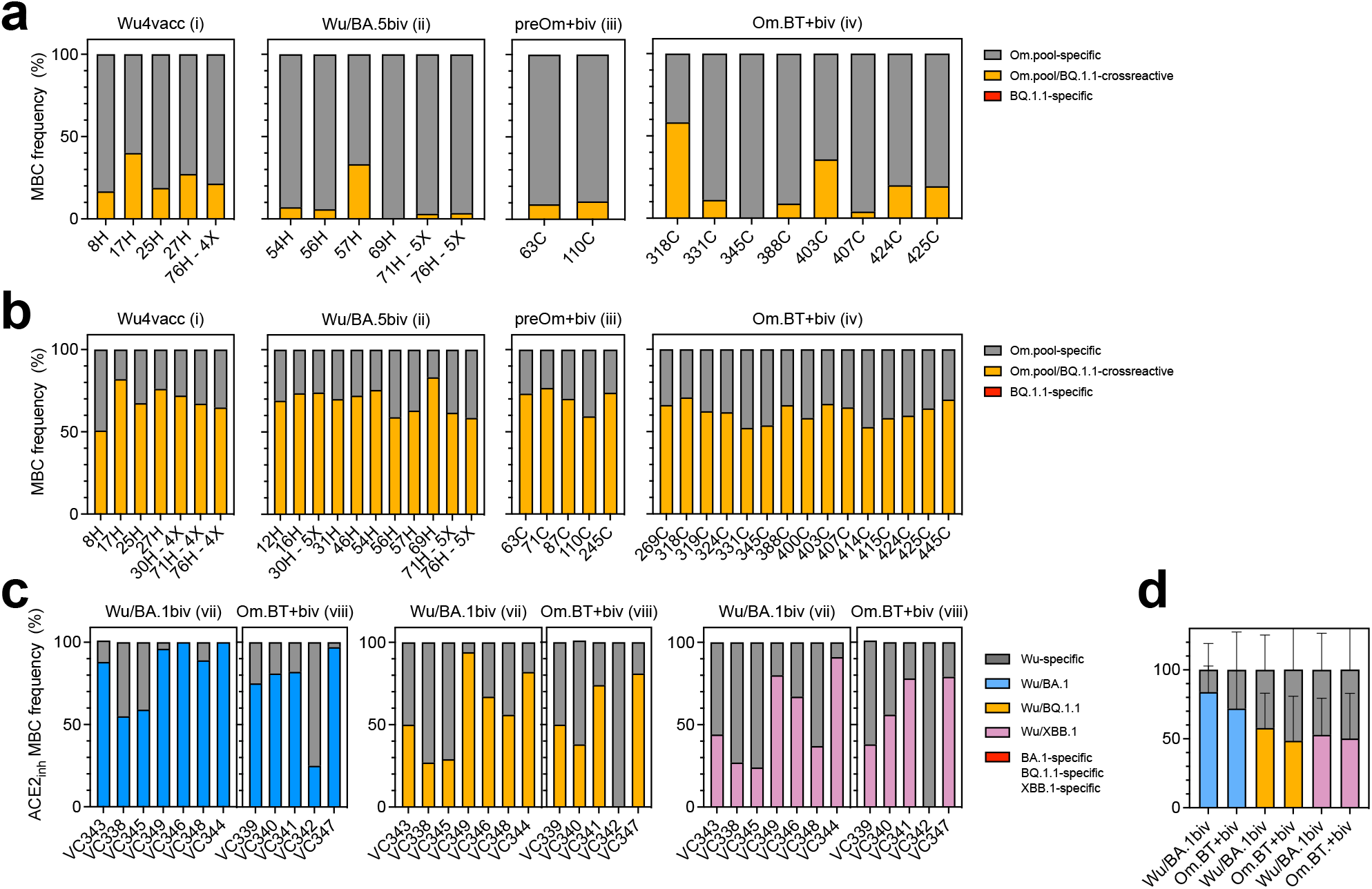
Subanalysis of cross-reactivity of vaccine-elicited MBCs. **a-c**, Analysis of cross-reactivity with BQ.1.1 of Omicron-specific (a) and Wu/Omicron-cross-reactive (b). Om.pool, MBCs reactive to Omicron BA.1, BA.2 or BA.5 RBDs in cohorts i-iv. **c, d**, Individual (c) and mean±sd (d) frequencies of Wu-G614-specific (grey), Omicron-specific (red) and Wu/Omicron-cross-reactive (blue for BA.1, yellow for BQ.1.1 and purple for XBB.1) MBCs showing inhibition of binding of ACE2 to RBD from donors of cohorts vii and viisi.

**Supplementary Table 1.** Kinetics of monomeric human ACE2 binding to immobilized SARS-CoV-2 variant RBDs as measured by biolayer interferometry.

|  | **K<sub>D</sub> (nM)** | **k<sub>on</sub> (M<sup>-1</sup>s<sup>-1</sup>)** | **k<sub>off</sub>(s<sup>-1</sup>)** |
| --- | --- | --- | --- |
| **Wuhan-Hu-1** | 101.1 ± 7.3 | 1.30 x 10<sup>5</sup> | 1.31 x 10<sup>-2</sup> |
| **BA.4/5** | 12.8 ± 2.2 | 1.45 x 10<sup>5</sup> | 1.82 x 10<sup>-3</sup> |
| **BA.2.75.2** | 26.2 ± 1.7 | 1.35 x 10<sup>5</sup> | 3.52 x 10<sup>-3</sup> |
| **BQ.1.1** | 13.7 ± 1.4 | 1.39 x 10<sup>5</sup> | 1.89 x 10<sup>-3</sup> |
| **XBB.1** | 88.4 ± 11.9 | 1.47 x 10<sup>5</sup> | 1.29 x 10<sup>-2</sup> |

Values are presented as mean ± standard deviation

**Supplementary Table 2.** Kinetics of monomeric human ACE2 binding to immobilized SARS-CoV-2 variant RBDs as measured by Surface Plasmon Resonance.

| RBD | average<br>KD (nM) | stdev<br>(KD) (nM) | averaged<br>ka (1/Ms) | stdev(ka)<br>(1/Ms) | averaged<br>kd (1/s) | stdev(kd)<br>(1/s) | Number<br>of<br>replicates |
| --- | --- | --- | --- | --- | --- | --- | --- |
| Wu-Hu-WT | 117.93 | 7.86 | 6.09E+04 | 6.65E+03 | 6.09E+04 | 6.65E+03 | 3 |
| Wu-Hu<br>E340A | 110.25 | 4.35 | 5.42E+04 | 1.36E+04 | 5.95E-03 | 1.26E-03 | 3 |
| BA.2 | 32.40 | 2.90 | 2.27E+04 | 1.92E+03 | 7.34E-04 | 8.47E-05 | 3 |
| BA.4/5 | 28.72 | 1.28 | 2.69E+04 | 4.17E+02 | 7.73E-04 | 2.26E-05 | 2 |

Averaged binding affinity (KD in nM unit), corresponding standard deviation, and number of replicates for each RBD-ACE2 pair. N/A: standard deviation calculation not applicable because only one replicate produces data within the instrument's limit. NB: no binding; N/A: not applicable.

**Supplementary Table 3.** CryoEM data collection and refinement statistics.

|  | SARS-CoV-2 BQ.1.1 RBD -<br>ACE2-S309<br>PDB<br>EMD | SARS-CoV-2 XBB.1 RBD -<br>ACE2-S309<br>PDB<br>EMD |
| --- | --- | --- |
| **Data collection and processing** |  |  |
| Magnification | 105,000 | 105,000 |
| Voltage (kV) | 300 | 300 |
| Electron exposure (e<sup>-</sup>/Å<sup>2</sup>) | 60 | 60 |
| Defocus range (μm) | -0.5 - -2.5 | -0.2 - -3.0 |
| Pixel size (Å) | 0.843 | 0.843 |
| Symmetry imposed | C1 | C1 |
| Final particle images (no.) | 564,989 | 281,957 |
| Map resolution (Å) | 3.2 | 3.1 |
| FSC threshold | 0.143 | 0.143 |
| Map sharpening *B* factor (Å<sup>2</sup>) | -150 | -120 |
| **Validation** |  |  |
| MolProbity score | 0.95 | 1.51 |
| Clashscore | 1.07 | 5.94 |
| Poor rotamers (%) | 0.46 | 0.48 |
| Ramachandran plot |  |  |
| Favored (%) | 97.26 | 96.87 |
| Allowed (%) | 2.64 | 2.71 |
| Disallowed (%) | 0.1 | 0.42 |

**Supplementary Table 4.**
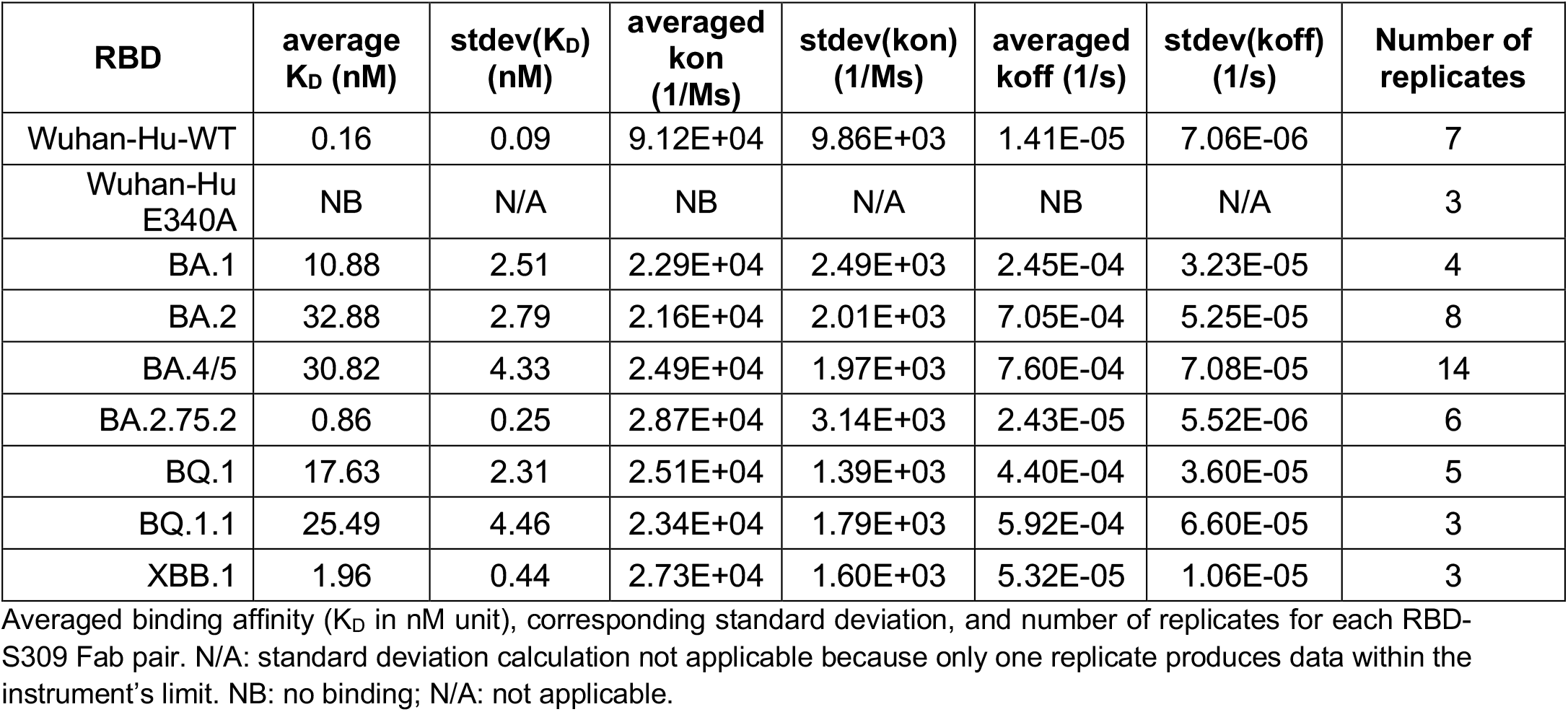
Kinetics of S309 Fab binding to immobilized SARS-CoV-2 variant RBDs as measured by Surface Plasmon Resonance.

| RBD | average<br>K<sub>D</sub> (nM) | stdev(K<sub>D</sub>)<br>(nM) | averaged<br>kon<br>(1/Ms) | stdev(kon)<br>(1/Ms) | averaged<br>koff (1/s) | stdev(koff)<br>(1/s) | Number of<br>replicates |
| --- | --- | --- | --- | --- | --- | --- | --- |
| Wuhan-Hu-WT | 0.16 | 0.09 | 9.12E+04 | 9.86E+03 | 1.41E-05 | 7.06E-06 | 7 |
| Wuhan-Hu<br>E340A | NB | N/A | NB | N/A | NB | N/A | 3 |
| BA.1 | 10.88 | 2.51 | 2.29E+04 | 2.49E+03 | 2.45E-04 | 3.23E-05 | 4 |
| BA.2 | 32.88 | 2.79 | 2.16E+04 | 2.01E+03 | 7.05E-04 | 5.25E-05 | 8 |
| BA.4/5 | 30.82 | 4.33 | 2.49E+04 | 1.97E+03 | 7.60E-04 | 7.08E-05 | 14 |
| BA.2.75.2 | 0.86 | 0.25 | 2.87E+04 | 3.14E+03 | 2.43E-05 | 5.52E-06 | 6 |
| BQ.1 | 17.63 | 2.31 | 2.51E+04 | 1.39E+03 | 4.40E-04 | 3.60E-05 | 5 |
| BQ.1.1 | 25.49 | 4.46 | 2.34E+04 | 1.79E+03 | 5.92E-04 | 6.60E-05 | 3 |
| XBB.1 | 1.96 | 0.44 | 2.73E+04 | 1.60E+03 | 5.32E-05 | 1.06E-05 | 3 |

Averaged binding affinity (K<sub>D</sub> in nM unit), corresponding standard deviation, and number of replicates for each RBD-S309 Fab pair. N/A: standard deviation calculation not applicable because only one replicate produces data within the instrument's limit. NB: no binding; N/A: not applicable.

**Supplementary Table 5.**
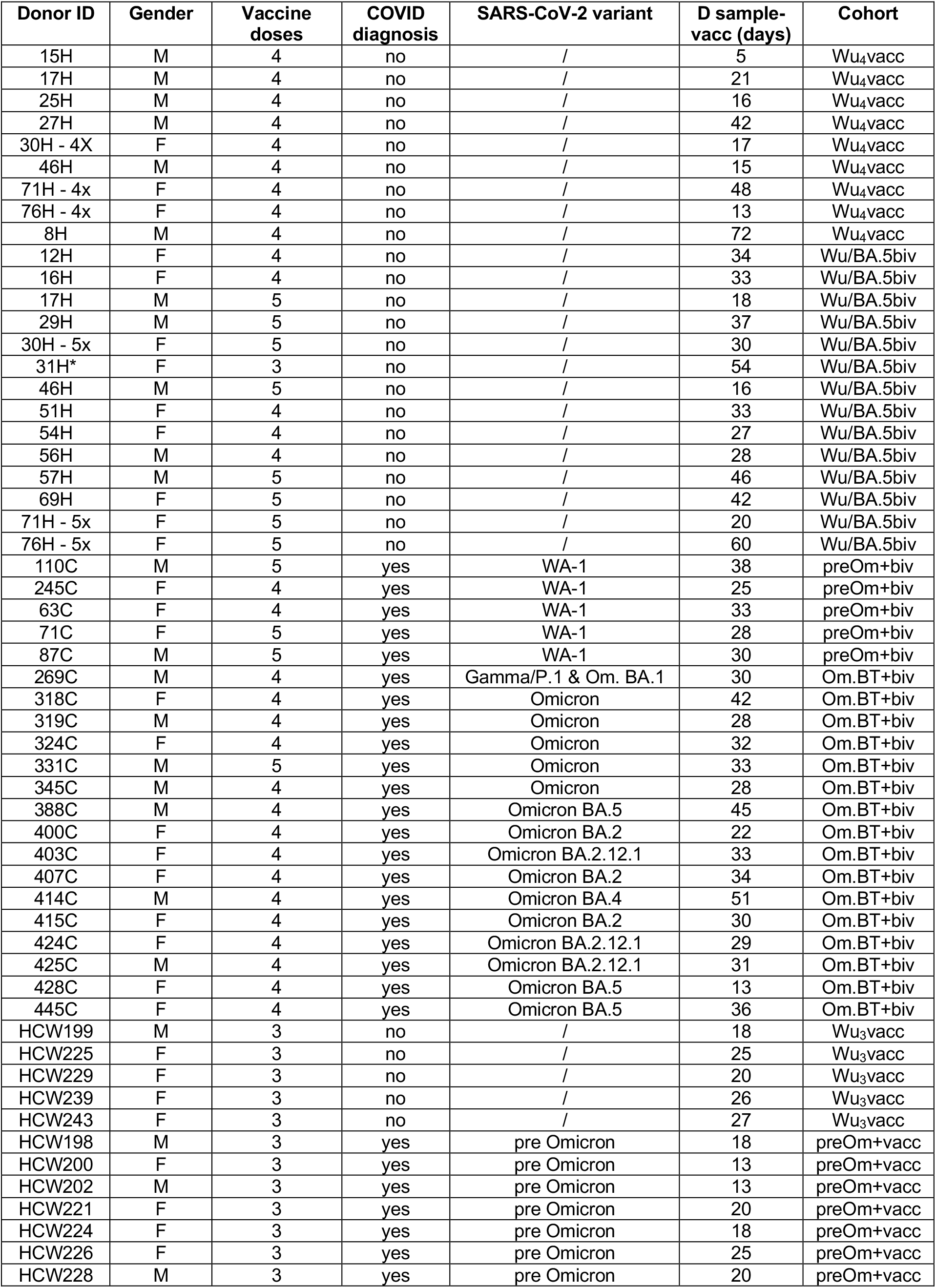
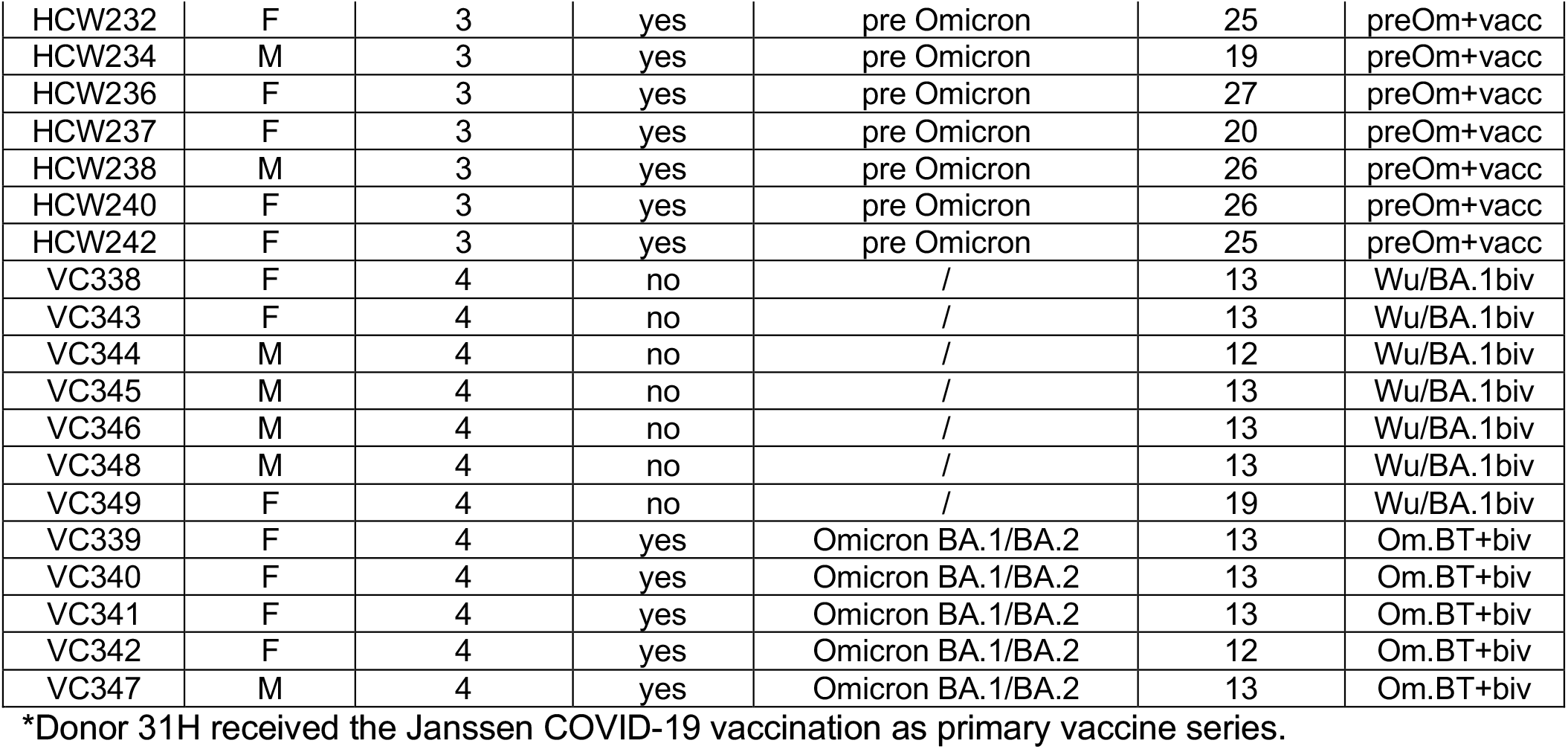
Donors’ demographics.

| Donor ID | Gender | Vaccine doses | COVID diagnosis | SARS-CoV-2 variant | D sample-vacc (days) | Cohort |
| --- | --- | --- | --- | --- | --- | --- |
| 15H | M | 4 | no | / | 5 | Wu<sub>4</sub>vacc |
| 17H | M | 4 | no | / | 21 | Wu<sub>4</sub>vacc |
| 25H | M | 4 | no | / | 16 | Wu<sub>4</sub>vacc |
| 27H | M | 4 | no | / | 42 | Wu<sub>4</sub>vacc |
| 30H - 4X | F | 4 | no | / | 17 | Wu<sub>4</sub>vacc |
| 46H | M | 4 | no | / | 15 | Wu<sub>4</sub>vacc |
| 71H - 4x | F | 4 | no | / | 48 | Wu<sub>4</sub>vacc |
| 76H - 4x | F | 4 | no | / | 13 | Wu<sub>4</sub>vacc |
| 8H | M | 4 | no | / | 72 | Wu<sub>4</sub>vacc |
| 12H | F | 4 | no | / | 34 | Wu/BA.5biv |
| 16H | F | 4 | no | / | 33 | Wu/BA.5biv |
| 17H | M | 5 | no | / | 18 | Wu/BA.5biv |
| 29H | M | 5 | no | / | 37 | Wu/BA.5biv |
| 30H - 5x | F | 5 | no | / | 30 | Wu/BA.5biv |
| 31H* | F | 3 | no | / | 54 | Wu/BA.5biv |
| 46H | M | 5 | no | / | 16 | Wu/BA.5biv |
| 51H | F | 4 | no | / | 33 | Wu/BA.5biv |
| 54H | F | 4 | no | / | 27 | Wu/BA.5biv |
| 56H | M | 4 | no | / | 28 | Wu/BA.5biv |
| 57H | M | 5 | no | / | 46 | Wu/BA.5biv |
| 69H | F | 5 | no | / | 42 | Wu/BA.5biv |
| 71H - 5x | F | 5 | no | / | 20 | Wu/BA.5biv |
| 76H - 5x | F | 5 | no | / | 60 | Wu/BA.5biv |
| 110C | M | 5 | yes | WA-1 | 38 | preOm+biv |
| 245C | F | 4 | yes | WA-1 | 25 | preOm+biv |
| 63C | F | 4 | yes | WA-1 | 33 | preOm+biv |
| 71C | F | 5 | yes | WA-1 | 28 | preOm+biv |
| 87C | M | 5 | yes | WA-1 | 30 | preOm+biv |
| 269C | M | 4 | yes | Gamma/P.1 & Om. BA.1 | 30 | Om.BT+biv |
| 318C | F | 4 | yes | Omicron | 42 | Om.BT+biv |
| 319C | M | 4 | yes | Omicron | 28 | Om.BT+biv |
| 324C | F | 4 | yes | Omicron | 32 | Om.BT+biv |
| 331C | M | 5 | yes | Omicron | 33 | Om.BT+biv |
| 345C | M | 4 | yes | Omicron | 28 | Om.BT+biv |
| 388C | M | 4 | yes | Omicron BA.5 | 45 | Om.BT+biv |
| 400C | F | 4 | yes | Omicron BA.2 | 22 | Om.BT+biv |
| 403C | F | 4 | yes | Omicron BA.2.12.1 | 33 | Om.BT+biv |
| 407C | F | 4 | yes | Omicron BA.2 | 34 | Om.BT+biv |
| 414C | M | 4 | yes | Omicron BA.4 | 51 | Om.BT+biv |
| 415C | F | 4 | yes | Omicron BA.2 | 30 | Om.BT+biv |
| 424C | F | 4 | yes | Omicron BA.2.12.1 | 29 | Om.BT+biv |
| 425C | M | 4 | yes | Omicron BA.2.12.1 | 31 | Om.BT+biv |
| 428C | F | 4 | yes | Omicron BA.5 | 13 | Om.BT+biv |
| 445C | F | 4 | yes | Omicron BA.5 | 36 | Om.BT+biv |
| HCW199 | M | 3 | no | / | 18 | Wu<sub>3</sub>vacc |
| HCW225 | F | 3 | no | / | 25 | Wu<sub>3</sub>vacc |
| HCW229 | F | 3 | no | / | 20 | Wu<sub>3</sub>vacc |
| HCW239 | F | 3 | no | / | 26 | Wu<sub>3</sub>vacc |
| HCW243 | F | 3 | no | / | 27 | Wu<sub>3</sub>vacc |
| HCW198 | M | 3 | yes | pre Omicron | 18 | preOm+vacc |
| HCW200 | F | 3 | yes | pre Omicron | 13 | preOm+vacc |
| HCW202 | M | 3 | yes | pre Omicron | 13 | preOm+vacc |
| HCW221 | F | 3 | yes | pre Omicron | 20 | preOm+vacc |
| HCW224 | F | 3 | yes | pre Omicron | 18 | preOm+vacc |
| HCW226 | F | 3 | yes | pre Omicron | 25 | preOm+vacc |
| HCW228 | M | 3 | yes | pre Omicron | 20 | preOm+vacc |
| HCW232 | F | 3 | yes | pre Omicron | 25 | preOm+vacc |
| HCW234 | M | 3 | yes | pre Omicron | 19 | preOm+vacc |
| HCW236 | F | 3 | yes | pre Omicron | 27 | preOm+vacc |
| HCW237 | F | 3 | yes | pre Omicron | 20 | preOm+vacc |
| HCW238 | M | 3 | yes | pre Omicron | 26 | preOm+vacc |
| HCW240 | F | 3 | yes | pre Omicron | 26 | preOm+vacc |
| HCW242 | F | 3 | yes | pre Omicron | 25 | preOm+vacc |
| VC338 | F | 4 | no | / | 13 | Wu/BA.1biv |
| VC343 | F | 4 | no | / | 13 | Wu/BA.1biv |
| VC344 | M | 4 | no | / | 12 | Wu/BA.1biv |
| VC345 | M | 4 | no | / | 13 | Wu/BA.1biv |
| VC346 | M | 4 | no | / | 13 | Wu/BA.1biv |
| VC348 | M | 4 | no | / | 13 | Wu/BA.1biv |
| VC349 | F | 4 | no | / | 19 | Wu/BA.1biv |
| VC339 | F | 4 | yes | Omicron BA.1/BA.2 | 13 | Om.BT+biv |
| VC340 | F | 4 | yes | Omicron BA.1/BA.2 | 13 | Om.BT+biv |
| VC341 | F | 4 | yes | Omicron BA.1/BA.2 | 13 | Om.BT+biv |
| VC342 | F | 4 | yes | Omicron BA.1/BA.2 | 12 | Om.BT+biv |
| VC347 | M | 4 | yes | Omicron BA.1/BA.2 | 13 | Om.BT+biv |

\*Donor 31H received the Janssen COVID-19 vaccination as primary vaccine series.

**Supplementary Table 6.** Kidney transplant recipients’ and healthcare workers’ demographics.

| Donor ID | Gender | No. of immunosuppressive drugs* | Vaccine doses | COVID diagnosis | SARS-CoV-2 variant | D sample-vaccine (days) | Cohort |
| --- | --- | --- | --- | --- | --- | --- | --- |
| KTR-004 | M | 2 | 4 | no | / | 83 | KTR Wu<sub>4</sub>vacc |
| KTR-007 | M | 3 | 4 | no | / | 140 | KTR Wu<sub>4</sub>vacc |
| KTR-009 | M | 2 | 4 | no | / | 46 | KTR Wu<sub>4</sub>vacc |
| KTR-010 | F | 3 | 4 | no | / | 51 | KTR Wu<sub>4</sub>vacc |
| KTR-011 | M | 2 | 4 | no | / | 80 | KTR Wu<sub>4</sub>vacc |
| KTR-013 | F | 1 | 4 | no | / | 92 | KTR Wu<sub>4</sub>vacc |
| KTR-026 | M | 2 | 4 | no | / | 63 | KTR Wu<sub>4</sub>vacc |
| KTR-027 | F | 3 | 4 | no | / | 35 | KTR Wu<sub>4</sub>vacc |
| KTR-030 | M | 2 | 4 | no | / | 71 | KTR Wu<sub>4</sub>vacc |
| KTR-039 | M | 2 | 4 | no | / | 77 | KTR Wu<sub>4</sub>vacc |
| KTR-042 | M | 3 | 4 | no | / | 40 | KTR Wu<sub>4</sub>vacc |
| KTR-047 | M | 2 | 4 | no | / | 76 | KTR Wu<sub>4</sub>vacc |
| KTR-050 | M | 2 | 4 | no | / | 84 | KTR Wu<sub>4</sub>vacc |
| KTR-054 | M | 2 | 4 | no | / | 18 | KTR Wu<sub>4</sub>vacc |
| KTR-056 | M | 2 | 4 | no | / | 70 | KTR Wu<sub>4</sub>vacc |
| KTR-059 | M | 2 | 4 | no | / | 78 | KTR Wu<sub>4</sub>vacc |
| KTR-060 | M | 2 | 4 | no | / | 71 | KTR Wu<sub>4</sub>vacc |
| KTR-061 | M | 2 | 4 | no | / | 54 | KTR Wu<sub>4</sub>vacc |
| KTR-071 | F | 3 | 4 | no | / | 35 | KTR Wu<sub>4</sub>vacc |
| KTR-083 | M | 3 | 4 | no | / | 82 | KTR Wu<sub>4</sub>vacc |
| KTR-085 | F | 2 | 4 | no | / | 71 | KTR Wu<sub>4</sub>vacc |
| KTR-094 | M | 2 | 4 | no | / | 41 | KTR Wu<sub>4</sub>vacc |
| KTR-095 | M | 3 | 4 | no | / | 74 | KTR Wu<sub>4</sub>vacc |
| KTR-096 | M | 2 | 4 | no | / | 43 | KTR Wu<sub>4</sub>vacc |
| KTR-101 | M | 2 | 4 | no | / | 72 | KTR Wu<sub>4</sub>vacc |
| KTR-102 | M | 3 | 4 | no | / | 133 | KTR Wu<sub>4</sub>vacc |
| KTR-017 | F | 2 | 4 | yes | pre Omicron | 58 | KTR preOm+Wu<sub>4</sub>vacc |
| KTR-021 | M | 2 | 4 | yes | pre Omicron | 84 | KTR preOm+Wu<sub>4</sub>vacc |
| KTR-031 | F | 2 | 4 | yes | pre Omicron | 59 | KTR preOm+Wu<sub>4</sub>vacc |
| KTR-084 | F | 3 | 4 | yes | pre Omicron | 65 | KTR preOm+Wu<sub>4</sub>vacc |
| KTR-099 | M | 2 | 4 | yes | pre Omicron | 93 | KTR preOm+Wu<sub>4</sub>vacc |
| HCW-001 | M | / | 3 | no | / | 90 | HCW Wu<sub>3</sub>vacc |
| HCW-002 | F | / | 3 | no | / | 78 | HCW Wu<sub>3</sub>vacc |
| HCW-003 | F | / | 3 | no | / | 76 | HCW Wu<sub>3</sub>vacc |
| HCW-004 | M | / | 3 | no | / | 90 | HCW Wu<sub>3</sub>vacc |
| HCW-005 | F | / | 3 | no | / | 83 | HCW Wu<sub>3</sub>vacc |
| HCW-008 | F | / | 3 | no | / | 86 | HCW Wu<sub>3</sub>vacc |
| HCW-009 | F | / | 3 | no | / | 81 | HCW Wu<sub>3</sub>vacc |
| HCW-011 | M | / | 3 | no | / | 93 | HCW Wu<sub>3</sub>vacc |
| HCW-012 | F | / | 3 | no | / | 94 | HCW Wu<sub>3</sub>vacc |
| HCW-013 | F | / | 3 | no | / | 55 | HCW Wu<sub>3</sub>vacc |
| HCW-016 | F | / | 3 | no | / | 6 | HCW Wu<sub>3</sub>vacc |
| HCW-017 | F | / | 3 | no | / | 50 | HCW Wu<sub>3</sub>vacc |
| HCW-018 | M | / | 3 | no | / | 110 | HCW Wu<sub>3</sub>vacc |
| HCW-019 | F | / | 3 | no | / | 29 | HCW Wu<sub>3</sub>vacc |
| HCW-020 | F | / | 3 | no | / | 90 | HCW Wu<sub>3</sub>vacc |
| HCW-021 | M | / | 3 | no | / | 28 | HCW Wu<sub>3</sub>vacc |
| HCW-022 | F | / | 3 | no | / | 28 | HCW Wu<sub>3</sub>vacc |
| HCW-023 | F | / | 3 | no | / | 22 | HCW Wu<sub>3</sub>vacc |
| HCW-024 | M | / | 3 | no | / | 35 | HCW Wu<sub>3</sub>vacc |
| HCW-025 | F | / | 3 | no | / | 94 | HCW Wu<sub>3</sub>vacc |
| HCW-026 | F | / | 3 | no | / | 28 | HCW Wu<sub>3</sub>vacc |
| HCW-027 | F | / | 3 | no | / | 38 | HCW Wu<sub>3</sub>vacc |
| HCW-028 | F | / | 3 | no | / | 29 | HCW Wu<sub>3</sub>vacc |
| HCW-031 | F | / | 3 | no | / | 35 | HCW Wu<sub>3</sub>vacc |
| HCW-033 | F | / | 3 | no | / | 31 | HCW Wu<sub>3</sub>vacc |
| HCW-037 | F | / | 3 | no | / | 45 | HCW Wu<sub>3</sub>vacc |
| HCW-038 | F | / | 3 | no | / | 61 | HCW Wu<sub>3</sub>vacc |
| HCW-039 | F | / | 3 | no | / | 23 | HCW Wu<sub>3</sub>vacc |
| HCW-010 | F | / | 3 | yes | pre Omicron | 69 | HCW preOm+Wu<sub>3</sub>vacc |
| HCW-014 | F | / | 3 | yes | pre Omicron | 89 | HCW preOm+Wu<sub>3</sub>vacc |
| HCW-015 | F | / | 3 | yes | pre Omicron | 86 | HCW preOm+Wu<sub>3</sub>vacc |
| HCW-029 | F | / | 3 | yes | pre Omicron | 77 | HCW preOm+Wu<sub>3</sub>vacc |
| HCW-030 | F | / | 3 | yes | pre Omicron | 35 | HCW preOm+Wu<sub>3</sub>vacc |
| HCW-034 | F | / | 3 | yes | pre Omicron | 28 | HCW preOm+Wu<sub>3</sub>vacc |
| HCW-036 | F | / | 3 | yes | pre Omicron | 34 | HCW preOm+Wu<sub>3</sub>vacc |

\*any of the following: cyclosporin, tacrolimus, MMF/MPA, azathioprine, everolimus/sirolimus, belatacept or glucocorticoids

